# MAP Kinase OsMEK2 and OsMPK1 Signaling for Ferroptotic Cell Death in Rice-*Magnaporthe oryzae* Interactions

**DOI:** 10.1101/2020.06.26.174292

**Authors:** Sarmina Dangol, Raksha Singh, Khoa Nam Nguyen, Yafei Chen, Juan Wang, Hyeon Gu Lee, Byung Kook Hwang, Nam-Soo Jwa

## Abstract

Mitogen-activated protein kinase (MAPK) signaling is required for plant cell death responses to invading microbial pathogens. Ferric ions and reactive oxygen species (ROS) accumulate in rice (*Oryza sativa*) tissues undergoing cell death during *Magnaporthe oryzae* infection. Here, we report that rice MAP kinase (OsMEK2 and OsMPK1) signaling cascades are involved in iron- and ROS-dependent ferroptotic cell death responses of rice to *M. oryzae* infection. OsMEK2 interacted with OsMPK1 in the cytoplasm, and OsMPK1 moved from the cytoplasm into the nucleus to bind to the OsWRKY90 transcription factor. *OsMEK2* expression may trigger OsMPK1-OsWRKY90 signaling pathways in the nucleus. Avirulent *M. oryzae* infection in *ΔOsmek2* mutant rice did not trigger iron and ROS accumulation and lipid peroxidation, and also downregulated *OsMPK1*, *OsWRKY90*, *OsRbohB*, and *OsPR-1b* expression. However, *OsMEK2* overexpression induced ROS-and iron-dependent cell death in rice during *M. oryzae* infection. The downstream MAP kinase (*OsMPK1*) overexpression induced ROS- and iron-dependent ferroptotic cell death in the compatible rice-*M*. *oryzae* interaction. These data suggest that the OsMEK2-OsMPK1-OsWRKY90 signaling cascade is involved in the ferroptotic cell death in rice. The small-molecule inducer erastin triggered iron- and lipid ROS-dependent, but *OsMEK2*-independent, ferroptotic cell death in *ΔOsmek2* mutant plants during *M. oryzae* infection. Disease-related cell death was lipid ROS-dependent and iron-independent in the *ΔOsmek2* mutant plants. These combined results suggest that *OsMEK2* and *OsMPK1* expression positively regulates iron- and ROS-dependent ferroptotic cell death via OsMEK2-*OsMPK1*-*OsWRKY90* signaling pathways, and blast disease (susceptibility)-related cell death was ROS-dependent but iron-independent in rice-*M. oryzae* interactions.

## INTRODUCTION

Plants have evolved effective innate immune system responses to avert the invasion of microbial pathogens in their natural habitat (Dodds et al., 2010; Schwessinger et al., 2012; Fu and Dong, 2013). Plant immune system responses are mediated by pattern-triggered immunity (PTI) and effector-triggered immunity (ETI), which are effectively upregulated inside plant cells in response to pathogen infection (Jones and Dangl, 2006). PTI is activated by plant perception of conserved microbial structures, called pathogen-associated molecular patterns (PAMPs), via the transmembrane pattern recognition receptors (PRRs) (Zipfel, 2008). ETI is activated by plant recognition of specific pathogen effector molecules via intracellular nucleotide-binding leucine-rich repeat (NLR) receptors, called resistance (R) proteins (Jones and Dangl, 2006; Block and Alfano, 2011; Oh and Martin, 2011). The two immune systems trigger a series of molecular signaling events that lead to diverse cellular responses including transcriptional reprogramming, synthesis of defense-related proteins, reactive oxygen species (ROS) burst, and iron- and ROS-dependent ferroptotic cell death (Boller and He, 2009; Dangol et al., 2019).

Mitogen-activated protein (MAP) kinase (MAPK) signaling pathways have pivotal roles in plant defense, immunity, and hypersensitive cell death responses to pathogen attack (Ishihama et al., 2011; Meng and Zhang, 2013; Thulasi Devendrakumar et al., 2018). However, the downstream signaling networks activated by defense-related MAPKs have not been completely defined in plants. Plant MAPK cascades proceed through three central kinases: MAPK kinase kinase (MAPKKK); MAPK kinase (MAPKK), also known as <u>M</u>APK and <u>E</u>RK (extracellular signal–regulated kinase) <u>k</u>inase (MEK); and MAP kinase (MAPK or MPK) (Meng and Zhang, 2013). These kinases are sequentially phosphorylated as MAPKKKs activate downstream MAPKKs (MEKs), which subsequently activate MAPKs (Rodriguez et al., 2010). Phosphorylation of MAPKs may promote their nuclear translocation to target other kinases, proteins, or transcription factors in the nucleus (Khokhlatchev et al., 1998; Rodriguez et al., 2010). MAPKs can activate transcription factors such as WRKYs. The Arabidopsis genome encodes 60 MAPKKKs, 10 MAPKKs, and 20 MAPKs (Ichimura et al., 2002). A previous study showed that Arabidopsis innate immune responses are mediated by a MAP kinase signaling cascade (MEKK1, MKK4/MKK5, and MPK3/MPK6) and WRKY22/WRKY29 transcription factors (Asai et al., 2002). Arabidopsis MPK3 and MPK6 are involved in ETI (Tsuda et al., 2009; Meng and Zhang, 2013; Su et al., 2018). Pathogen-responsive MAPK cascades (MEKK1-MKK4/MKK5-MPK3/MPK6 and MEKK1-MKK1/2-MPK4) have pivotal roles in defense signaling against pathogen attack in *Arabidopsis thaliana* (Pitzschke, 2009; Rasmussen et al., 2012; Meng and Zhang, 2013). Nearly two decades ago, Yang et al. (2001) identified a tobacco MAPKK (NtMEK2) upstream of both salicylic acid-induced protein kinase (SIPK) and wounding-induced protein kinase (WIPK). Expression of a constitutively active *NtMEK2* mutant induced hypersensitive response (HR)-like cell death and defense responses in tobacco. Many kinases in MAPK cascades, including MAPKKK, MEK, SIPK/WIPK, and MAPK, are involved in *N* gene-mediated resistance to tobacco mosaic virus in tobacco (Jin et al., 2002, 2003; Liu et al., 2004). Tobacco WRKY/MYB transcription factors downstream of MAPK cascades have crucial roles in regulating *N*-mediated resistance to TMV (Liu et al., 2004). Silencing of *MEK2* (*SlMKK2*), *SlMPK2*, and *SlMKK4* in tomato disrupted the resistance to infection by *Xanthomonas campestris* pv. *vesicatoria* (*Xcv*) and *Botrytis cinerea* (Melech-Bonfil and Sessa, 2011; Li et al., 2014).

The rice genome contains 74 MAPKKK, 8 MAPKK, and 17 MAPK genes (Hamel et al., 2006; Reyna and Yang, 2006; Rao et al., 2010; Yang et al., 2015). We previously identified 74 nonredundant interactors with rice MAPKs and performed high-resolution mapping of the MAPK interactome network, which controls different signaling pathways underlying the cellular and physiological responses in rice (Singh et al., 2012). Rice MAP kinase kinase1 (OsMEK1) physically interacts with rice MAP kinase1 (OsMPK1), OsMPK6, and OsMPK5. OsMEK2 interacts with OsMPK1 and OsMPK6. OsMEK6 interacts with OsMPK1 and OsMPK5 (Singh et. al., 2012). Rice MAP kinase kinases (OsMAP2Ks or OsMEKs) may regulate multiple signaling pathways affecting many biological processes by associating with different sets of rice MAPK interactomes (Singh and Jwa, 2013). However, few kinase components in rice MAPK cascades are involved in immunity and defense responses in rice-pathogen interactions (Singh and Jwa, 2013; Yang et al., 2015). OsMKK10-2–mediated activation of OsMPK6 via specific phosphorylation subsequently induced *WRKY45* expression and blast (*Magnaporthe oryzae*) resistance in rice plants (Ueno et al., 2015). OsMPKK10-2 is involved in disease resistance and drought tolerance (Ma et al., 2017), physically interacts with OsMPK6 and OsMPK3, and phosphorylates the two OsMPKs, leading to *X. oryzae* pv. *oryzicola* (*Xoo*) resistance and drought tolerance (Ma et al., 2017). The MAP kinase module OsMKK3-OsMPK7-OsWRKY30 is involved in induced resistance to *Xoo* infection in rice (Jalmi and Sinha, 2016).

Cell death is a fundamental biological process that occurs during development, senescence, immunity, and stress resistance in multicellular organisms. Increases in ROS accumulation levels, including superoxide (·O_2_^−^), hydrogen peroxide (H_2_O_2_), and hydroxyl radical (·OH), occur during pathogen infection and are involved in basal immune responses, NLR-mediated hypersensitive cell death, and disease-associated cell death in plants (Greenberg and Yao, 2004; Choi et al., 2012, 2013; Jwa and Hwang, 2017). Ferroptosis differs from apoptosis, necrosis, and autophagy, and was first discovered in mammalian cells as a form of nonapoptotic iron-dependent cell death (Dixon et al., 2012). Ferroptotic cell death requires the accumulation of ROS, iron, and lipid peroxides (Stockwell et al., 2017). Iron homeostasis and ROS burst have important roles in activating defense responses against plant pathogens (Liu et al., 2007; Aznar et al., 2015). Powdery mildew (*Blumeria graminis* f.sp. *tritici*) infection in wheat (*Triticum aestivum*) epidermal cells induces ferric ion (Fe^3+^) accumulation at fungal attack sites to mediate the ROS (H_2_O_2_) burst (Liu et al., 2007). Heat shock stress induces ferroptosis-like cell death in Arabidopsis roots (Distéfano et al., 2017). Heat shock treatment of Arabidopsis root hairs at 55°C leads to iron and ROS accumulation, and triggers ferroptotic cell death.

We recently reported that iron- and ROS-dependent ferroptosis occurs in incompatible rice–*Magnaporthe oryzae* interactions (Dangol et al., 2019). This was the first plant pathosystem in which ferroptotic cell death was discovered (Caseys, 2019). Avirulent *M. oryzae* infection triggers iron and ROS (H_2_O_2_) accumulation at the cell death sites in rice tissues (Dangol et al., 2019). Highly reactive Fe^2+^ reacts with H_2_O_2_ to produce Fe^3+^ and ∙OH (Fenton, 1894; Pierre and Fontecave, 1999), which results in iron-dependent accumulation of toxic lipid ROS (Dixon et al., 2012). Iron is required for lipid peroxide accumulation. Iron and ROS accumulation and lipid peroxidation are blocked by the iron chelator deferoxamine, the lipophilic antioxidant ferrostatin-1, the actin polymerization inhibitor cytochalasin E, and the NADPH-oxidase inhibitor diphenyleneiodonium (DPI), thereby restricting HR cell death in rice (Dangol et al., 2019). By contrast, the RAS-selective lethal small molecule inducer erastin triggered iron-dependent ROS accumulation and glutathione depletion, which ultimately promoted virulent *M. oryzae-*induced ferroptotic cell death. Rice NADP-malic enzyme (NADP-ME) and NADPH-oxidase (Rboh) are ROS sources that have been proposed to be involved in iron- and ROS-dependent ferroptotic cell death (Singh et al., 2016; Dangol et al., 2019).

Our previous study identified rice mitogen-activated protein (MAP) kinase kinase 2 (OsMEK2) as a rice MAP interactor (Singh et al., 2012). Rice MAP kinase (OsMPK1) is an interactor of OsMEK2 and actively involved in *M. oryzae* infection (Singh et al., 2012; Ueno et al., 2015). Here, we use *OsMEK2* and *OsMPK1* to investigate whether rice MAPKs are involved in the signaling network that mediates ferroptotic cell death in rice–*M. oryzae* interactions. *OsMEK2* deletion via T-DNA insertion in the resistant rice cultivar Dongjin (DJ) suppressed iron- and ROS-dependent ferroptotic cell death, which ultimately induced susceptible responses to avirulent *M. oryzae* 007 infection. However, overexpression of OsMEK2 (35S:*OsMEK2*) in rice DJ induced iron- and ROS-dependent ferroptotic cell death against avirulent *M. oryzae* 007 infection. Treatment of the *OsMEK2*-deleted DJ (*ΔOsmek2*) mutant with the RAS-selective lethal small molecule inducer erastin induced the ROS burst and iron accumulation, which caused ferroptotic cell death in response to avirulent *M. oryzae* infection. Disease (susceptibility)-related cell death at the late stage [96 h post-inoculation (hpi)] of *M. oryzae* infection in the susceptible *ΔOsmek2* mutant is ROS-dependent and iron-independent. During *M. oryzae* infection, *OsMEK2* deletion and overexpression differentially regulated the expression of *OsMPK1*, *OsMPK6*, and the *OsWRKY90* transcription factor in the rice MAPK signaling pathways. *OsMPK1* overexpression in susceptible rice cultivar Nipponbarre (NB) induced iron- and ROS-mediated ferroptotic cell death against virulent *M. oryzae* PO6-6. These combined results indicate that *OsMEK2* and *OsMPK1* expression via MAPK signaling pathway positively regulates iron- and ROS-dependent ferroptotic cell death and plant immunity to *M. oryzae* infection.

## RESULTS

### Identification of the Rice MAPK Interactor OsMEK2

Our previous study isolated rice Mitogen-activated protein kinase kinases (MAPKKs or MEKs) that interacted with OsMAPKs using a rice leaf cDNA library and yeast two-hybrid analysis (Singh et al., 2012). Amino acid sequences of the isolated rice MEKs were aligned with those of Arabidopsis AtMAPKKs, and subsequently categorized into Groups A–D (Ichimura et al., 2002) (Fig. 1A). The plant MAPKKs contained 11 conserved subdomains (Supplemental Fig. S1). All of the aligned MAPKKs contained the active site domain [D(I/L/V)KP] and the conserved motif (S/T-X_5_-S/T, where X represents any amino acid residue) (Fig. 1A; Supplemental Fig. S1). The rice MAPKKs OsMEK1 and OsMEK2 belonged to Group A serine (S)/threonine (T) kinases. OsMEK1 amino acid sequence shared 54% and 53% homology with AtMKK1 and AtMKK2, respectively, whereas OsMEK2 had 62% sequence homology with both AtMKK1 and AtMKK2. A phylogenetic tree was generated using the neighbor-joining method of the Molecular Evolutionary Genetics Analysis Version 7.0 (MEGA7) to compare OsMEKs with Arabidopsis MAPKKs (Kumar et al., 2016) (Fig. 1B). Further phylogenetic analyses of rice MAPKs with pathogen-responsive MAPKs of Arabidopsis, tomato, and tobacco (*Nicotiana benthamiana*) indicated that OsMEK2 also shared 65–66% homology with tobacco NbMEK2 and tomato SlMKK1 (Supplemental Fig. S2). However, OsMEK2 has low levels of identity (37–39%) with tobacco NtMEK2 and tomato SlMKK2 (Supplemental Fig. S1). Based on the sequence alignment data of rice MAPKKs, OsMEK2 was selected to investigate whether rice MAPKKs are required for ferroptotic cell death signaling during rice–*M. oryzae* interactions.

**Figure 1.**
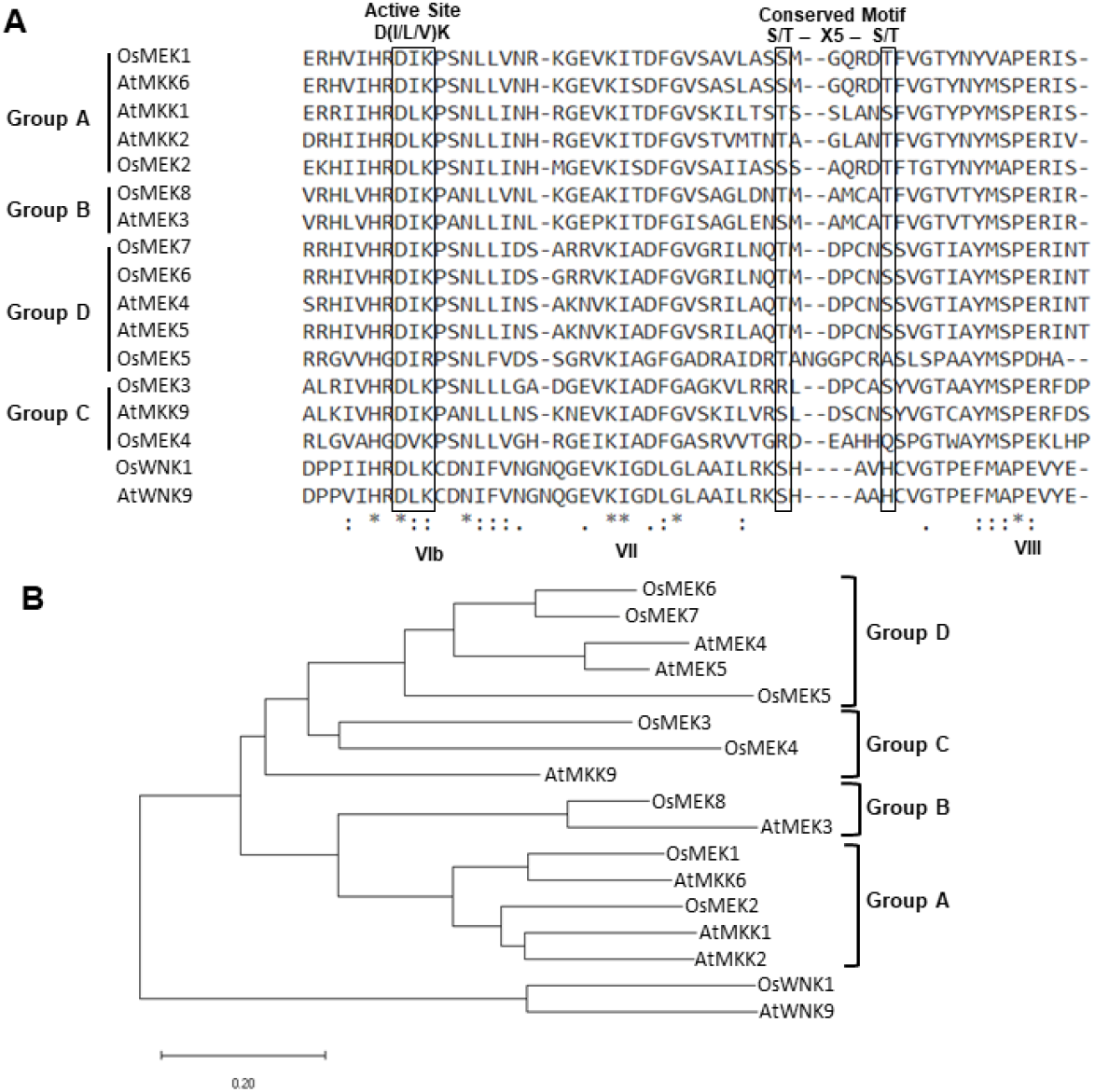
Amino acid sequence alignment and phylogenetic tree of rice MAPKKs (OsMEKs) with Arabidopsis MAPKKs. A, Amino acid sequence alignment of OsMEKs with Arabidopsis MAPKKs. Rice MAPKKs are aligned with Arabidopsis MAPKKs, which are categorized into four groups (Group A–D) using Clustal Omega (EMBL-EBI). The MAPKK active site [D(I/L/V)K] and conserved domain [S/T-X-S/T] are located between kinase subdomains VII and VIII (shown in yellow). B, Phylogenetic tree of OsMEKs with Arabidopsis MAPKKs was constructed using the neighbor-joining method based on Molecular Evolutionary Genetics Analysis Version 7.0 (MEGA7) (Kumar et al., 2016). Accession numbers of the plant MAPKKs are OsMEK1 (Os01g32660), OsMEK2 (Os06g05520), OsMEK3 (Os03g12390), OsMEK4 (Os02g46760), OsMEK5 (Os06g09190), OsMEK6 (Os02g54600), OsMEK7 (Os06g09180), OsMEK8 (Os06g27890), OsWNK1 (Os07g38530), AtMKK1 (At4g26070), AtMKK2 (At4g29810), AtMEK3 (NP_198860), AtMEK4 (At1g51660), AtMEK5 (At3g21220), AtMKK6 (At5g56580), AtMKK9 (At1g73500), and AtWNK9 (At3g04910).

*OsMEK2* was deleted in the wild-type (WT) rice cultivar DJ by T-DNA insertion mutagenesis (Jeon et al., 2000), and *ΔOsmek2* T-DNA insertion mutant seeds were obtained from the Rice Functional Genomic Express Database (RiceGE, Salk Institute Genomic Analysis Laboratory). The *OsMEK2* genomic DNA sequence contains nine exons and eight introns (Supplemental Fig. S3). The T-DNA insertion in *ΔOsmek2* was identified in the intronic region between the sixth and seventh exons (Fig. 2A). The genotypes of *ΔOsmek2* (M5) progeny were analyzed with primer sets LP+RP and LP+LB to detect transgene and homo/hetero selection, respectively (Fig. 2A). The *ΔOsmek2-2 and ΔOsmek2-4* mutants (M5) were identified as T-DNA insertion homozygous plants that lacked the specific PCR products.

**Figure 2.**
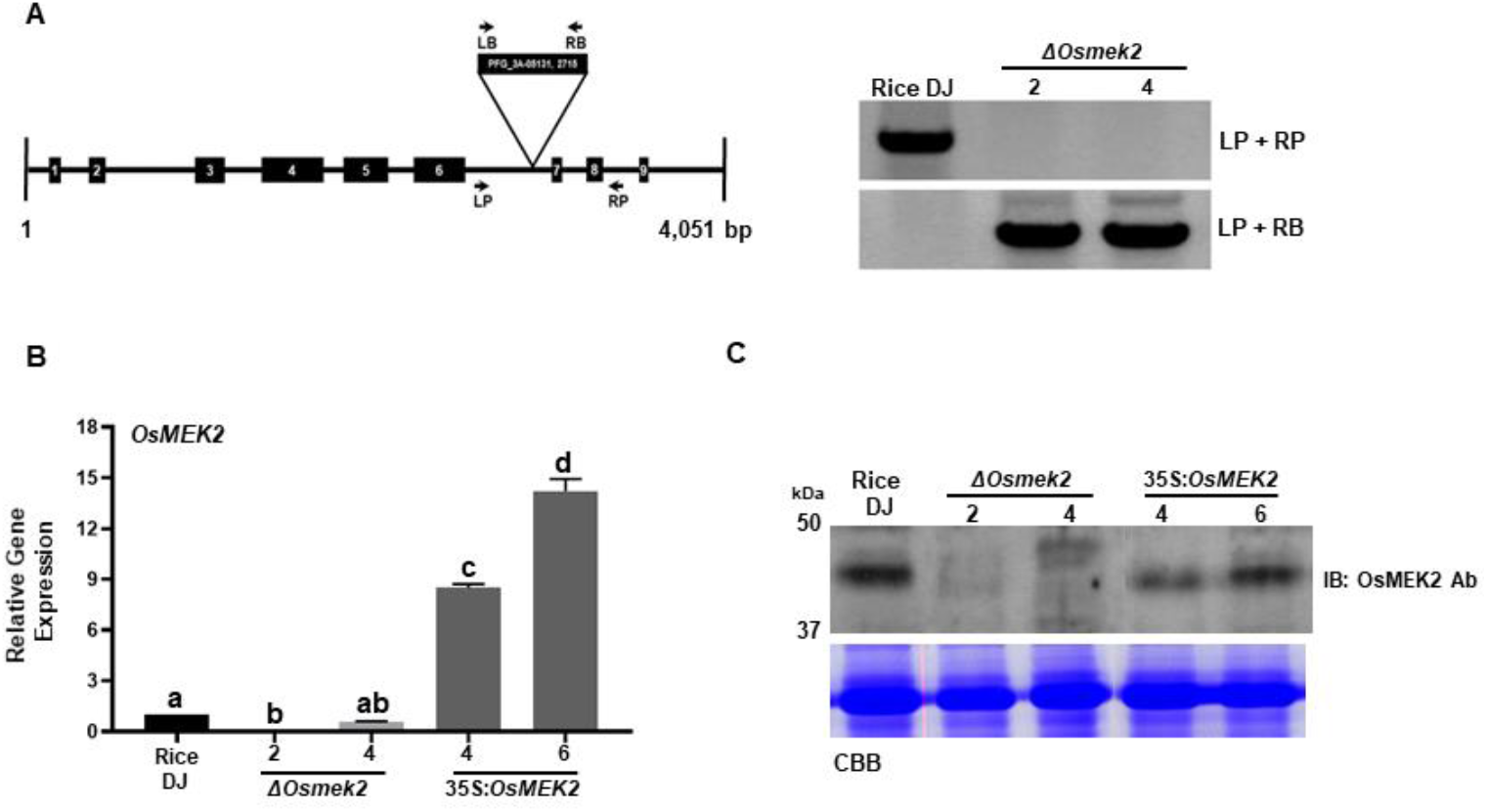
Genotyping, transcriptional, and immunoblotting analyses of *ΔOsmek2* and 35S:*OsMEK2* mutants. A, Genotyping of *ΔOsmek2* mutants. The schematic diagram shows the T-DNA insertion site in the *OsMEK2* gene. Exons and introns are depicted by solid boxes and lines, respectively. The T-DNA insertion *ΔOsmek2* mutant plants (M5) were detected using the gene primers (LP+RP) and the vector primers (LP+RB). LP, left primer; RP, right primer; LB, left border; RB, right border. B, Transcriptional analysis of *OsMEK2* expression in wild-type (WT) rice (cultivar DJ), *ΔOsmek2,* and 35S:*OsMEK2* mutants using RT-PCR and qRT-PCR. C, SDS-PAGE and immunoblotting assays of *OsMEK2* expression in wild-type (WT) rice (cultivar DJ), *ΔOsmek2* and 35S:*OsMEK2* mutant plants using OsMEK2 Ab (EnoGene^®^ E580135-A-SE) (~39 kDa). Ab, antibody; IB, immunoblot; PAGE, polyacrylamide gel electrophoresis.

35S:*OsMEK2* was overexpressed in Rice DJ under the control of CaMV 35S promoter. The T6 progeny of *OsMEK2* overexpression plants was selected to use in this study. *OsMEK2* expression in *ΔOsmek2* and 35S:*OsMEK2* mutant lines was examined by RT-PCR and immunoblotting with anti-OsMEK2 (Fig. 2, B and C). The RT-PCR and immunoblot analyses indicated that *OsMEK2* was not expressed in *ΔOsmek2-2 and ΔOsmek2-4* mutant lines, but distinctly expressed in 35S:*OsMEK2-4* and 35S:*OsMEK2-6* mutant lines, compared to the wild type Rice DJ. These combined data indicate that the *OsMEK2* gene is completely deleted in the selected *ΔOsmek2* mutant lines, but overexpressed in 35S:*OsMEK2* mutant lines. *ΔOsmek2-2* and 35S:*OsMEK2-4* were used in this study.

### *OsMEK2* Expression Patterns in Rice during *M. oryzae* Infection

We examined *OsMEK2* expression patterns in rice leaf sheaths of 5-week-old DJ rice plants by performing quantitative real-time RT-PCR analyses at different times after inoculation with virulent *M. oryzae* PO6-6 and avirulent *M. oryzae* 007 (Fig. 3). The results showed that avirulent *M. oryzae* 007 infection triggered the induction of *OsMEK2* expression at early infection times (1–12 hpi); *OsMEK2* expression was dramatically induced in rice leaf sheaths at 3 hpi, and then declined gradually by 48 hpi. By contrast, infection with virulent *M. oryzae* PO6-6 did not significantly induce *OsMEK2* expression by 48 hpi during compatible rice–*M. oryzae* interactions. Infection with avirulent *M. oryzae* 007 significantly reduced *OsMEK2* expression at late infection stages (72–96 hpi). These *OsMEK2* expression patterns (Fig. 3) suggest that early induction of *OsMEK2* expression in response to avirulent *M. oryzae* 007 infection is involved in rice defense signaling triggered by blast infection during the incompatible rice–*M. oryzae* interaction.

**Figure 3.**
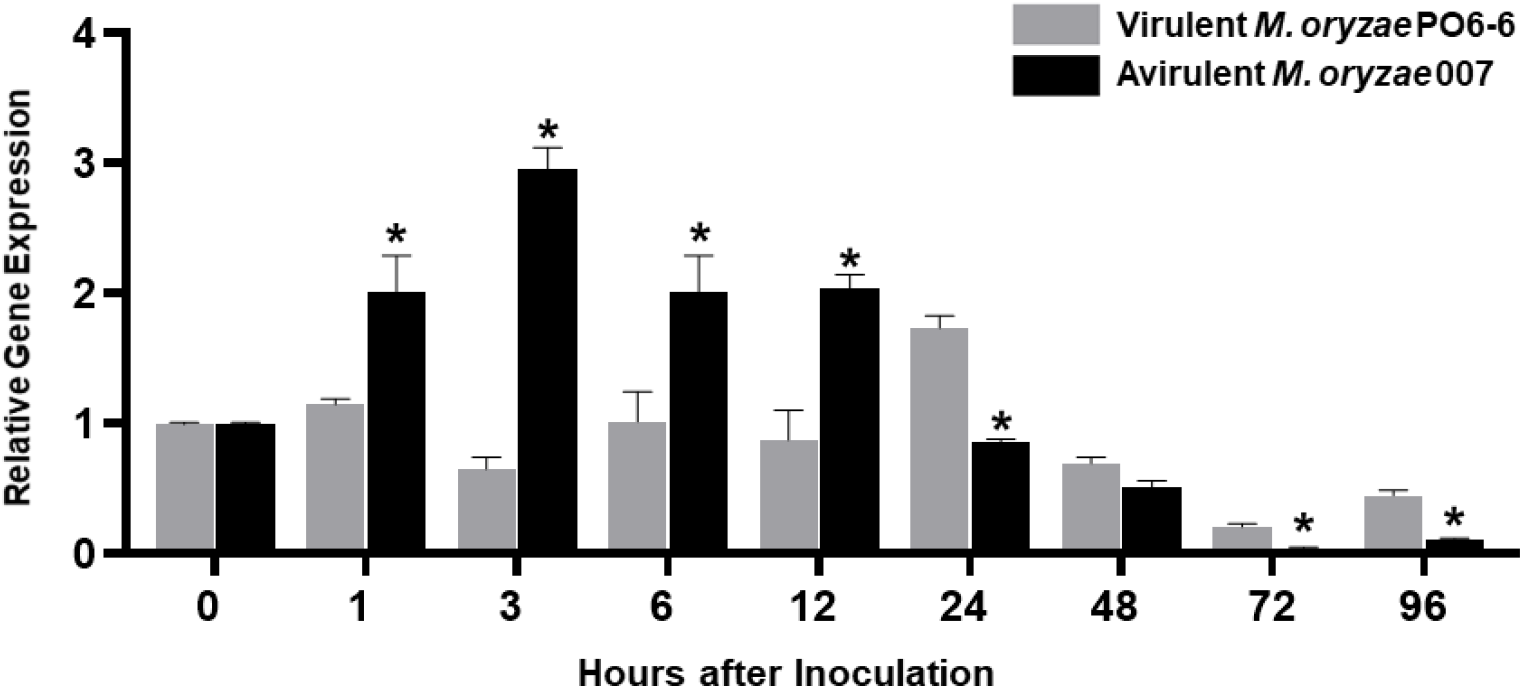
Quantitative real-time RT-PCR analysis of time-course expression of *OsMEK2* in rice leaf sheaths in the compatible and incompatible interactions of rice with *Magnaporthe oryzae*. Rice leaf sheaths were sampled at different time points after inoculation with virulent and avirulent *M. oryzae* PO6-6 and 007, respectively. *OsMEK2* expression was analyzed by quantitative RT-PCR. Relative gene expression of *OsMEK2* at each time point was calculated by normalizing with respect to expression of the internal control *OsUbiquitin* gene. Data represent the mean ± SD from three independent experiments. Asterisks above the columns indicate significant differences as analyzed by Student’s *t*-test (*P* < 0.05). hpi, hours post-inoculation.

### The *OsMEK2* Gene Is Required for Cell Death and Resistant Responses to *M. oryzae* Infection

To investigate whether rice MAPKKs are required for resistant responses to *M. oryzae* infection, we inoculated leaf sheaths and whole leaves of DJ, *ΔOsmek2* and 35S:*OsMEK2* mutant rice plants with a conidial suspension (4 × 10^5^ conidia mL^−1^) of avirulent *M. oryzae* 007 (Fig. 4). The infected cells were observed at 48 hpi. During infection, *M. oryzae* 007 grew poorly and caused cell death responses in leaf sheath epidermal cells of rice DJ and 35S:*OsMEK2* mutant plants (Fig. 4, A and B). By contrast, the blast fungus grew well with plentiful invasive hyphae (IH) in the invaded *ΔOsmek2* leaf sheath cells. Most of the infected *ΔOsmek2* cells had viable *M. oryzae* 007 colonizing hyphae (Fig. 4A). By contrast, avirulent *M. oryzae* 007 infection induced significantly more hypersensitive cell death in rice DJ and 35S:*OsMEK2* leaf sheaths than in *ΔOsmek2* leaf sheaths (Fig. 4B). Whole-leaf disease phenotypes were observed at 5 days after inoculation with *M. oryzae* 007 (Fig. 4C). Rice DJ and 35S:*OsMEK2* leaves displayed a typical resistant reaction with slightly elongated, necrotic, and brownish spots. By contrast, *ΔOsmek2* mutant leaves displayed a typical susceptible reaction with large, elliptical, and grayish lesions (Fig. 4C). These combined results indicate that *OsMEK2* deletion in the resistant WT rice cultivar DJ rendered resistance ineffective and induced susceptibility (disease) in response to avirulent *M. oryzae* infection of *ΔOsmek2* mutant plants. However, *OsMEK2* overexpression in rice DJ enhances the cell death and resistance responses to rice blast disease.

**Figure 4.**
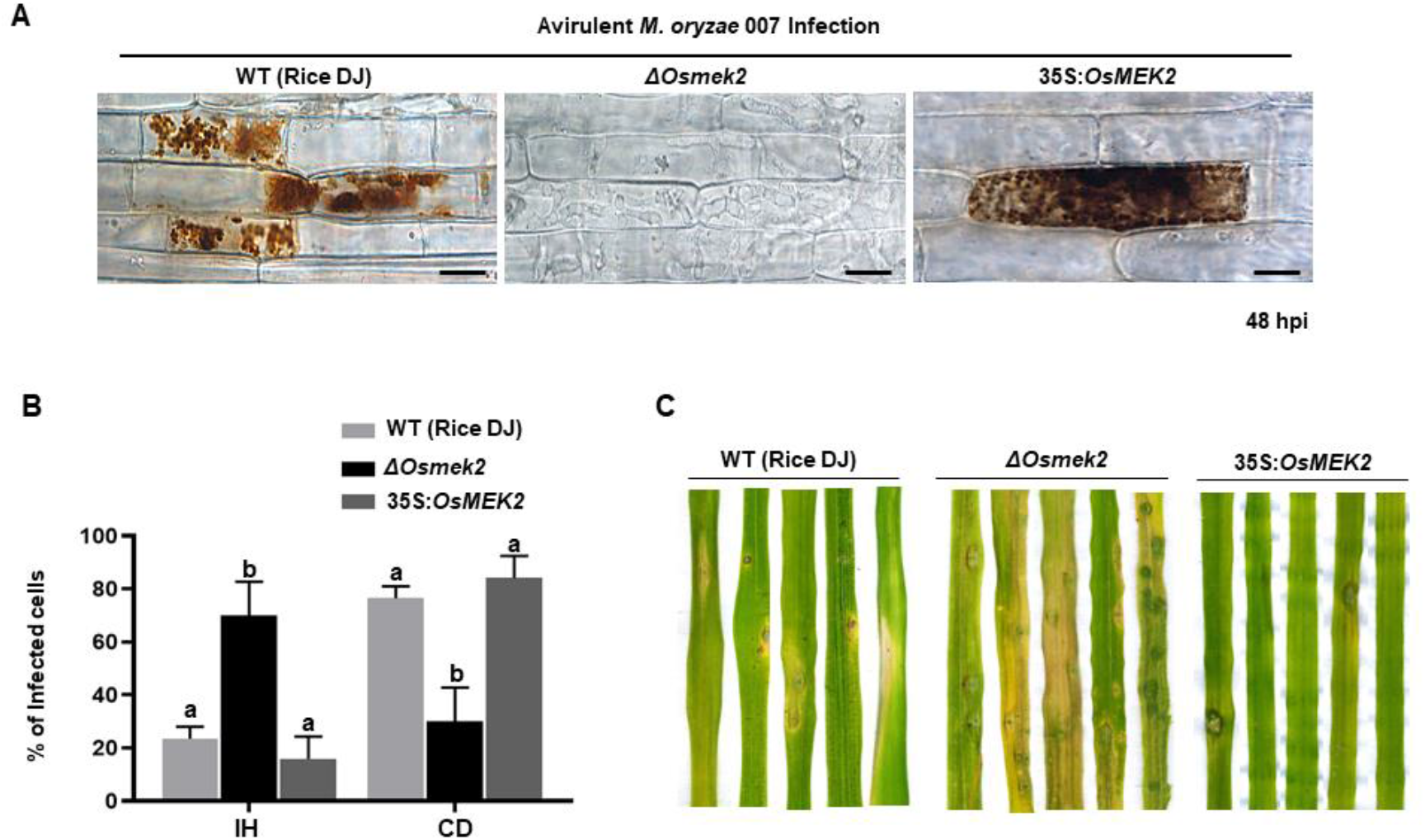
Avirulent *Magnaporthe oryzae* 007 infection causes susceptible responses in the *ΔOsmek2* mutant, but resistant responses in the wild-type rice and 35:*OsMEK2* mutant. A, Images of rice sheath epidermal cells infected with *M. oryzae* 007 (48 hpi). Rice leaf sheaths were inoculated with a conidial suspension (4×10^5^ conidia mL^−1^). *M. oryzae* 007 grew well and produced invasive hyphae in the *OsMEK2-*deleted mutant (*ΔOsmek2*) rice, but induced hypersensitive cell death in wild-type (WT) rice cultivar DJ and *OsMEK2-*overexpressed (35:*OsMEK2).*mutant plants. Images were captured using a fluorescence microscope. hpi, hours post-inoculation. Scale bars=20 μm. B, Quantification of cell death and invasive hyphae in rice sheath cells infected with *M. oryzae* 007 (48 hpi). Results are presented as mean values ± SD; *n*=4 leaf sheaths from different plants. Different letters above the bars indicate significantly different means (*P*<0.05) as analyzed by Fisher’s protected least significant difference (LSD) test. IH, invasive hyphae; CD, cell death. C, Disease types of rice leaves in wild-type rice (DJ), *ΔOsmek2* and 35:*OsMEK2* mutants. Two-week-old rice seedlings were spray-inoculated with a conidial suspension (4×10^5^ conidia mL^−1^) of *M. oryzae* 007. Diseased leaves were photographed at 5 days after inoculation. Disease types indicate a resistant-type lesion (slightly elongated, necrotic brownish spots) and a susceptible-type lesion (large, elliptical, grayish, and expanded lesions). Experiments were repeated three times with similar results.

### *OsMEK2* Deletion and Overexpression Differentially Regulates *OsMPK1*,*OsMPK6*, and *OsWRKY90* Expression during *M. oryzae* Infection

A previous study reported that rice OsMAP2K2 (OsMEK2) interacted with and phosphorylated OsMAPKs, such as OsMPK1 and OsMPK6 (Singh et al., 2012). Here, we analyzed *OsMPK1*, *OsMPK6*, and *OsWRKY90* expression in leaf sheaths of DJ, *ΔOsmek2* and 35S:*OsMEK2* mutant plants by performing real-time qRT-PCR at different time intervals after inoculation with avirulent *M. oryzae* 007 (Fig. 5). *OsMEK2* deletion in DJ plants distinctly downregulated *OsMPK1* expression throughout the course of *M. oryzae* infection. However, *OsMEK2* overexpression did not upregulate expression of *OsMPK1* and *OsMPK6* in DJ plants. By contrast, *OsMPK6* downregulation in *ΔOsmek2* leaf sheath cells was observed at early infection stages 3–12 hpi. *OsMPK6* (or *OsMPK1*) activation by *OsMKK10-2* is required for the induction of *OsWRKY45* expression and blast resistance in rice (Ueno et al., 2015). OsMPK1 is the pathogen-responsive MAPK that is involved in disease resistance (Fig. 5; Singh et al., 2012; Ueno et al., 2015). Bimolecular fluorescence complementation (BiFC) analysis in rice leaf sheath indicates that OsMPK1 physically interacts with the OsWRKY80 transcription factor (Singh et al., 2012) and subsequently OsWRKY90 (Shen et al., 2012) as its downstream target. In plant disease resistance networks, WRKY transcription factors can associate with MAPK cascades and regulate downstream defense-related genes in the nucleus (Pandey and Somssich, 2009; Ishihama et al., 2011; Jalmi and Sinha, 2016). Avirulent *M. oryzae* 007 infection significantly upregulated *OsWRKY90* expression in DJ and 35S:*OsMEK2* leaf sheaths, but did not affect *OsWRKY90* expression in *ΔOsmek2* leaf sheaths at all tested time points after inoculation (Fig. 5). This indicates that OsMEK2, the rice MAP2K, targets the OsWRKY90 transcription factor to function as a positive regulator of resistance to avirulent *M. oryzae* infection. Rice plant resistance to *M. oryzae* infection is markedly enhanced by overexpression of *OsWRKY45*, *OsWRKY53*, and *OsWRKY89* (Chujo et al., 2007; Shimono et al., 2007; Wang et al., 2007).

**Figure 5.**
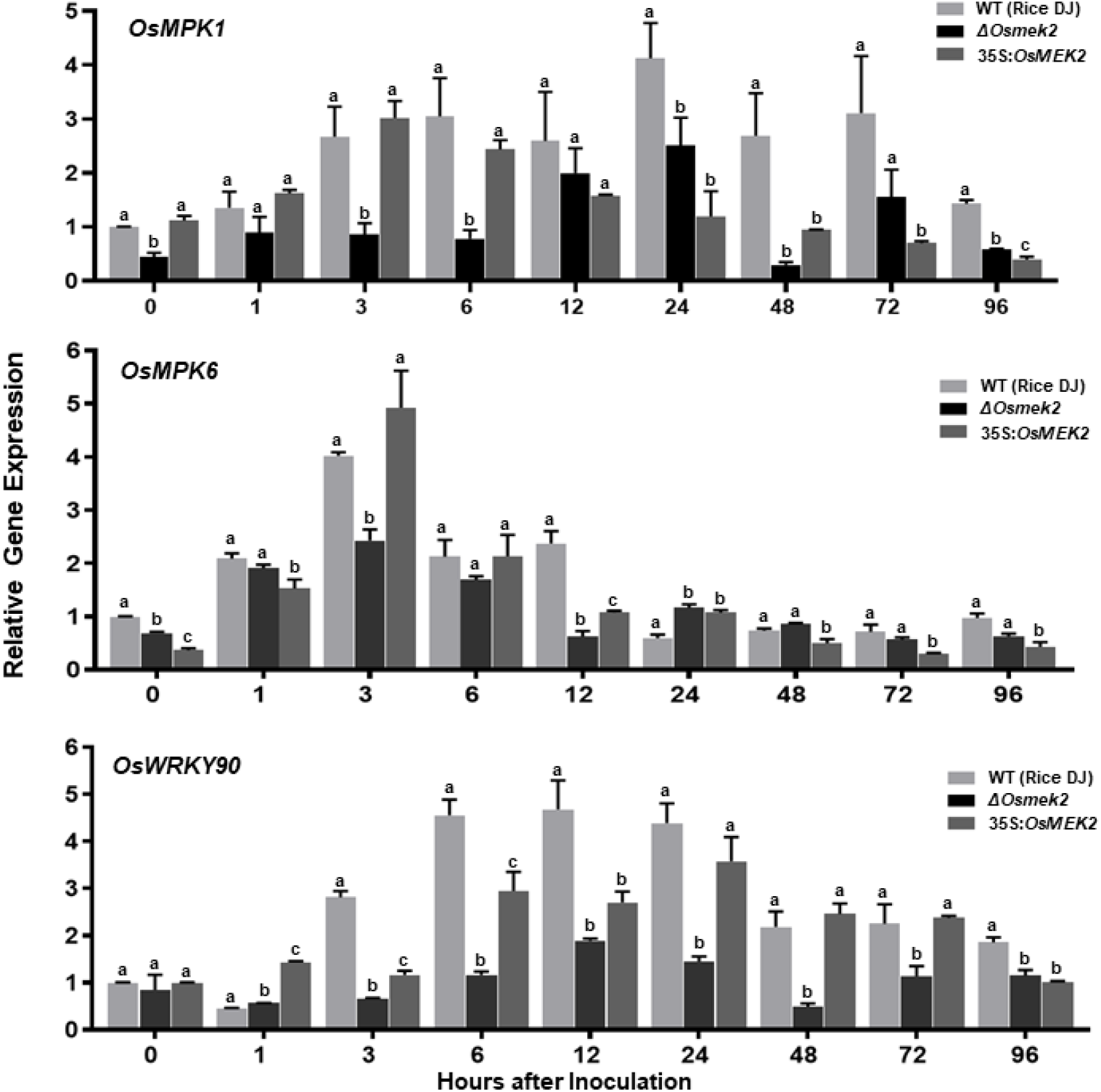
Quantitative real-time RT-PCR analysis of time-course expression of the *OsMEK2* interactors *OsMPK1*, *OsMPK6*, and *OsWRKY90* in leaf sheaths of wild-type (WT) rice (Cultivar DJ), *ΔOsmek2* and 35S:*OsMEK2* mutant plants infected with avirulent *Magnaporthe oryzae* 007. Leaf sheaths of wild-type (cultivar DJ), *ΔOsmek2* and 35S:*OsMEK2* mutant plants were sampled at different time points after inoculation, followed by total RNA extraction. Relative gene expression of *OsMPK1*, *OsMPK6*, and *OsWRKY90* at each time point was obtained by normalizing with respect to the expression of the internal control *OsUbiquitin* gene. Data represent the means ± SD from three independent experiments. Different letters above the bars indicate significantly different means (*P*<0.05) as analyzed by Fisher’s protected least significant difference (LSD) test.

### *OsMEK2* Expression Positively Regulates Expression of Defense-Related Genes *OsPR-1b* and *OsPAL1* in Rice during *M. oryzae* 007 Infection

We next investigated the expression patterns of some defense-related genes that are induced in response to pathogen infection in rice, such as pathogenesis-related protein 1b (*OsPR-1b*), phenylalanine ammonia lyase1 (*OsPAL1*), ascorbate peroxidase1 (*OsAPX1*), and *OsAPX2* (Nakashita et al., 2001; Agrawal et al., 2003; Xie et al., 2011). We quantified the expression levels of these defense-related genes in rice DJ, *ΔOsmek2* and 35S:*OsMEK2* mutant leaf sheaths by performing real-time qRT-PCR analysis during avirulent *M. oryzae* 007 infection (Supplemental Fig. S4). *OsPR-1b* expression was induced in rice DJ and 35S:*OsMEK2* at all tested times, whereas it was only induced in *ΔOsmek2* at 96 hpi. *OsPAL1* was distinctly induced in 35S:*OsMEK2* leaf sheaths during avirulent *M. oryzae* 007 infection. *OsPAL1* expression patterns did not significantly differ in rice DJ and *ΔOsmek2* leaf sheath cells at 12–96 hpi. *OsAPX1* and *OsAPX2* expression was gradually upregulated in *ΔOsmek2* mutant leaf sheath cells at 12–72 hpi (Supplemental Fig. S4). These results indicate that *OsMEK2* expression positively regulates *OsPR-1b* and *OsPAL1* expression in rice during avirulent *M. oryzae* 007 infection.

### *OsMEK2* Is Required for ROS and Ferric Ion Accumulation and Lipid Peroxidation in Rice-*M. oryzae* Interactions

We analyzed ROS and ferric ion (Fe^3+^) accumulation and lipid [malondialdehyde (MDA)] peroxidation in leaf sheath cells of rice DJ, *ΔOsmek2* and 35S:*OsMEK2* mutant plants during avirulent *M. oryzae* 007 infection to determine whether *OsMEK2* is involved in iron- and ROS-dependent ferroptotic cell death (Fig. 6).

**Figure 6.**
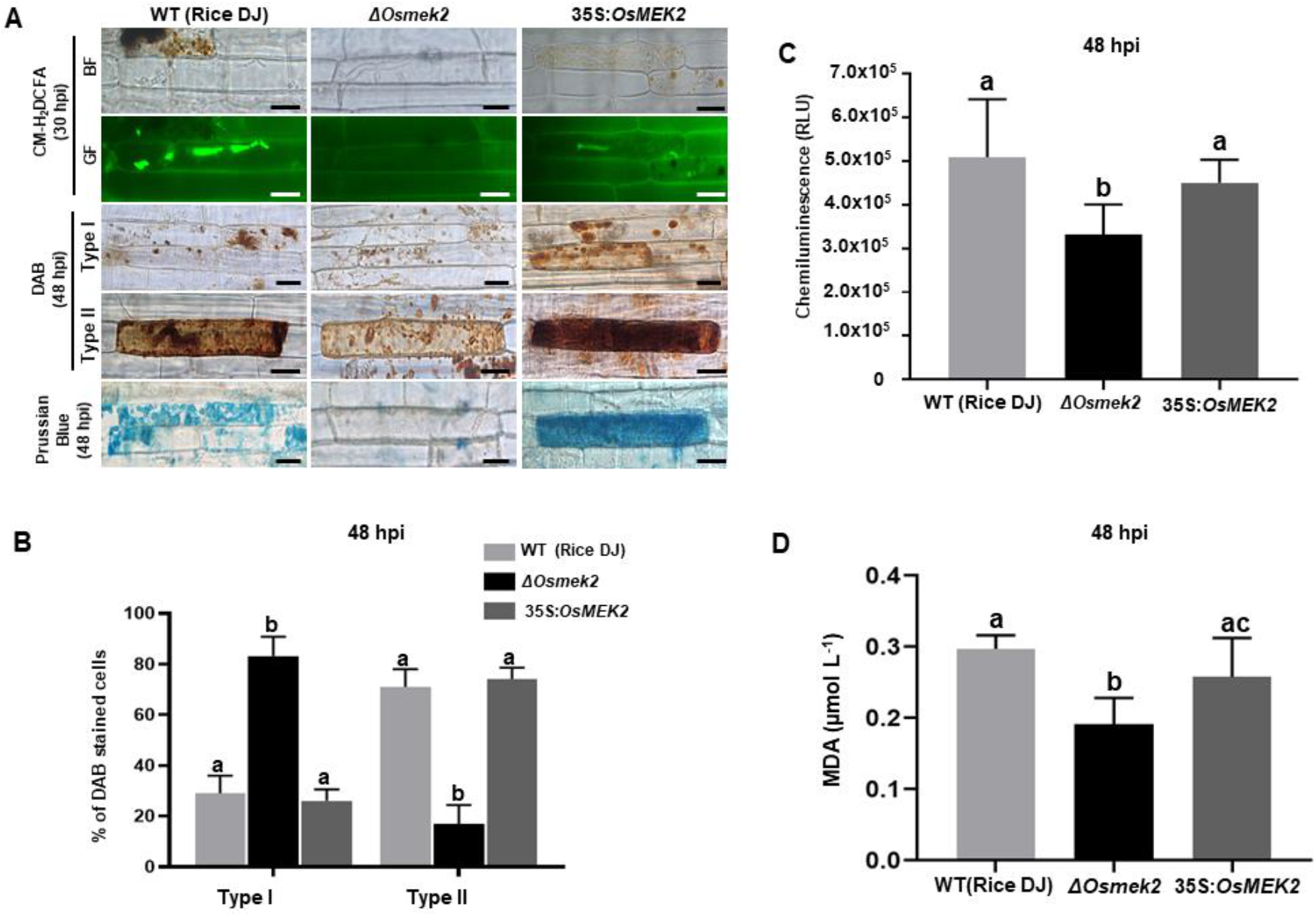
*OsMEK2* deletion and overexpression in wild-type rice DJ differentially regulates ROS and ferric Ion (Fe^3+^) accumulation and lipid peroxidation in leaf sheaths infected with avirulent *Magnaporthe oryzae* 007. A, CM-H_2_DCFDA (green fluorescence), DAB, and Prussian blue (blue color) staining shows accumulation of ROS (H_2_O_2_) and ferric ion (Fe^3+^) in rice leaf sheath epidermal cells of the wild-type (WT) rice cultivar DJ, *OsMEK2-*deleted (*ΔOsmek2*) and *OsMEK2-*overexpressed (35:*OsMEK2)* mutants during *M. oryzae* infection. Scale bar=20 μm. B, Quantification of DAB-stained cells at 48 h after inoculation. The DAB-stained cells were categorized into two phenotypes: Type I, infected cells that display no or weak DAB staining; and Type II, infected cells that display strong DAB staining. Results are presented as mean values ± SD; *n*=4 leaf sheaths from different plants. C, Quantification of ROS production in rice leaf sheaths at 48 h after inoculation. ROS production was quantified by a luminol-based assay using a GloMax^®^ 96 Microplate Luminometer (Promega). Values are means ± SD of total relative luminescent units (RLU) (*n*=10). D, Lipid (MDA) peroxidation (MDA) determination in rice leaf sheaths at 48 h after inoculation. Results are presented as mean values ± SD; *n*=4 leaf sheaths from different plants. Images were captured using a fluorescence microscope (Zeiss equipped with Axioplan 2) with bright field and a combination of excitation (450‒490 nm) and emission (515‒565 nm) GF filters. Experiments were repeated three times with similar results. Different letters above the bars indicate significantly different means (*P*<0.05) as analyzed by Fisher’s protected least significant difference (LSD) test. BF, bright field; GF, green fluorescence; hpi, hours post-inoculation; MDA, malondialdehyde.

ROS accumulation and localization in rice cells was detected with histochemical 5-(and 6-) chloromethyl-2′,7′-dichlorodihydrofluorescein diacetate acetyl ester (CM-H_2_DCFDA) and 3,3′-diaminobenzidine (DAB) staining (Fryer et al., 2002; Chi et al., 2009; Kristiansen et al., 2009). The ROS-sensitive CM-H_2_DCFDA dye is an indicator that can be used to monitor ROS localization in living plant cells (Kristiansen et al., 2009). CM-H_2_DCFDA (green fluorescence) and DAB (dark brown) staining revealed that ROS (H_2_O_2_) strongly accumulated inside and around invasive hyphae (IH) in rice DJ and 35S:*OsMEK2* epidermal cells at 36–48 hpi (Fig. 6A). By contrast, ROS did not accumulate around invasive hyphae (IH) in *ΔOsmek2* mutant epidermal cells after avirulent *M. oryzae* 007 infection. CM-H_2_DCFDA-specific ROS-localized fluorescence was clearly visible around invasive hyphae (IH) and cellular membranes in rice DJ and 35S:*OsMEK2* cells, whereas ROS-localized fluorescence was absent or weakly visible around invasive hyphae (IH) in *ΔOsmek2* mutant cells at 30 hpi (Fig. 6A). DAB is oxidized by H_2_O_2_ in the presence of peroxidase to generate a dark brown precipitate, which indicates the presence and distribution of H_2_O_2_ in plant cells (Fryer et al., 2002; Kristiansen et al., 2009). We classified DAB-stained cells into two phenotypes: Type I infected cells display no or weak DAB staining, and Type II infected cells display strong DAB staining (Fig. 6B). Most of the infected cells displayed strong brown staining (Type II phenotype) in rice DJ and 35S:*OsMEK2* cells, indicating relatively high levels of ROS accumulation in cells infected with avirulent *M. oryzae* 007. By contrast, significantly fewer *ΔOsmek2* mutant cells displayed DAB staining at 48 hpi. A chemiluminescent assay with a luminometer revealed that ROS levels were significantly lower in *ΔOsmek2* mutant cells than in DJ and 35S:*OsMEK2* cells at 48 hpi (Fig. 6A, B and C).

Ferric ion (Fe^3+^) accumulation and localization in rice cells was detected by Prussian blue (blue color) staining of rice DJ, *ΔOsmek2* and 35S:*OsMEK2* mutant leaf sheath cells at 48 hpi with avirulent *M. oryzae* 007 (Fig. 6A). The results showed that rice DJ and 35S:*OsMEK2* epidermal cells displayed strong blue staining, whereas *ΔOsmek2* mutant epidermal cells did not display blue stain. Next, we analyzed oxidative damage and lipid peroxidation in rice DJ, *ΔOsmek2* and 35S:*OsMEK2* mutant leaf sheath cells at 48 hpi with avirulent *M. oryzae* 007 (Fig. 6D) by performing the MDA assay as described previously (Zhang et al., 2009; Dangol et al., 2019). Lipid peroxidation levels were significantly lower in *ΔOsmek2* mutant cells than in rice DJ and 35S:*OsMEK2* cells (Fig. 6D), suggesting that reduced ROS and Fe^3+^ levels in *ΔOsmek2* cells also reduce oxidative stress levels. These combined results indicate that OsMEK2 has crucial roles in ROS and Fe^3+^ accumulation and lipid peroxidation during the ferroptotic cell death response in rice.

### Erastin Triggers Iron- and ROS-Dependent Ferroptotic Cell Death in *ΔOsmek2* Mutant Plants during *M. oryzae* Infection

Erastin is a small molecule inducer that triggers ferroptotic cell death in mammals and plants (Dixon et al., 2012; Dangol et al., 2019). Treatment with 10 μM erastin triggered ROS (H_2_O_2_) and Fe^3+^ accumulation and cell death response in *ΔOsmek2* mutant leaf sheaths during avirulent *M. oryzae* 007 infection (Fig. 7). However, mock (water) or 10 μM erastin treatment did not trigger ROS (H_2_O_2_) and Fe^3+^ accumulation in healthy rice DJ leaf sheaths (Supplemental Fig. S5). CM-H_2_DCFDA and DAB staining detected H_2_O_2_ accumulation in *ΔOsmek2* leaf sheath cells at 30–48 hpi with *M. oryzae* conidial suspension containing 10 μM erastin (Fig. 7, A and B). Erastin treatment during *M. oryzae* 007 infection induced H_2_O_2_ accumulation in *ΔOsmek2* mutant cells at 48 hpi as detected with a luminometer (Fig. 7D). Erastin induced Fe^3+^ accumulation and increased the number of Prussian blue-stained cells in *ΔOsmek2* leaf sheaths at 48 hpi (Fig. 7, A and C). Iron-dependent MDA peroxidation was upregulated at 48 hpi in *ΔOsmek2* mutant leaf sheath cells by treating with erastin (Fig. 7E). Erastin treatment significantly enhanced the cell death response in *ΔOsmek2* cells during *M. oryzae* infection compared to that in mock-treated *ΔOsmek2* cells during infection (Fig. 7, A and F). These combined results indicate that erastin triggers iron- and lipid ROS-dependent, but *OsMEK2*-independent, ferroptotic cell death in rice during *M. oryzae* infection.

**Figure 7.**
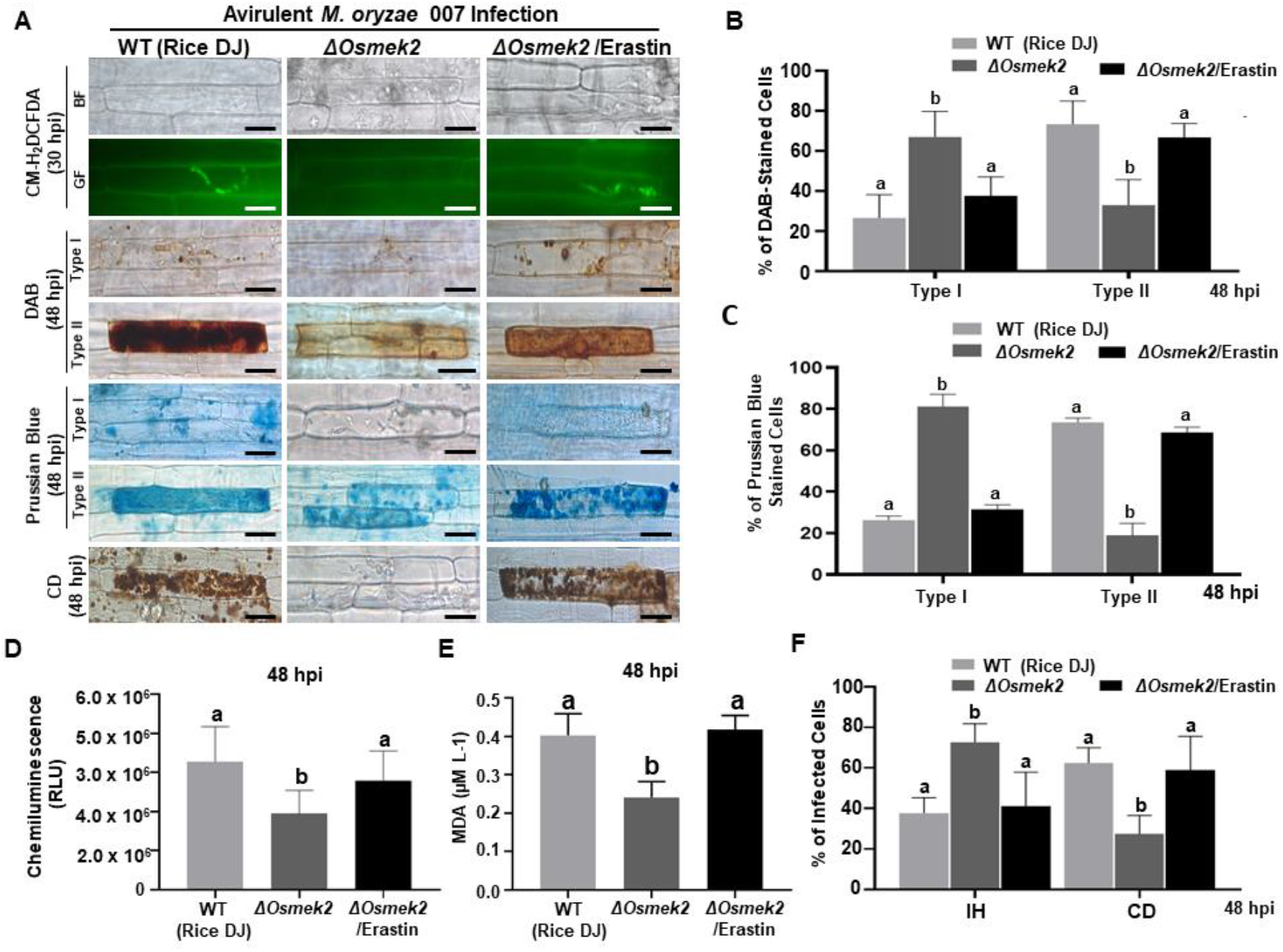
The small molecule inducer erastin triggers iron- and ROS-dependent ferroptotic cell death in the compatible *ΔOsmek2* mutant–*Magnaporthe oryzae* interaction. Leaf sheaths of WT rice DJ and *ΔOsmek2* mutant were inoculated with conidial suspensions (4×10^5^ conidia/mL) of *M. oryzae* 007 containing 10 μM erastin. A, Erastin treatment recovered ROS and ferric ion (Fe^3+^) accumulation and cell death in leaf sheaths of *ΔOsmek2* mutant during avirulent *M. oryzae* 007 infection. ROS accumulation in leaf sheath epidermal cells was detected by CM-H_2_DCFDA (green fluorescence) and DAB (dark brown color) staining. Prussian blue (blue color) staining shows ferric ion accumulation in rice cells. The images are representative of different leaf sheath samples from three independent experiments. Scale bars=20 μm. B, DAB-stained cell phenotypes. DAB-stained cells were divided into two phenotypes: Type I, cells that contain invasive hyphae (IH) but are weakly or not DAB-stained; and Type II, strongly DAB-stained cells with only a few poor hyphae. C, Quantification of Prussian blue-stained cells. Prussian blue-stained cells were divided into two phenotypes: Type I, cells that contain invasive hyphae (IH) but are weakly or not Prussian blue-stained; and Type II, strongly Prussian blue-stained with only a few poor hyphae. D, Quantification of ROS accumulation. ROS accumulation was monitored using a GloMax^®^ 96 Microplate Luminometer (Promega). Values are means ± SD of total relative luminescent units (RLU) (*n*=10). E, Determination of lipid (MDA) peroxidation in leaf sheaths at 48 h after inoculation. Results are presented as means ± SD; *n*=4 leaf sheaths from different plants. F, Quantification of infected cell phenotypes in rice leaf sheaths. Images were taken using a fluorescence microscope (Zeiss equipped with Axioplan 2) with bright field and green fluorescence (GF) filters. Experiments were repeated three times with similar results. Results are presented as means ± SD; *n*=4 leaf sheaths. Different letters above the bars indicate significantly different means (*P*<0.05) as analyzed by Fisher’s protected least significant difference (LSD) test. IH, invasive hyphae; CD, cell death; BF, bright field; GF, green fluorescence; hpi, hours post-inoculation.

### Disease-Related Cell Death Is ROS-Dependent but Iron-Independent in *ΔOsmek2* Mutant Plants during the Late Stage of *M. oryzae* Infection

ROS (H_2_O_2_) and Fe^3+^ did not accumulate in healthy rice DJ leaf sheaths at 72 and 92 h after treatment with 10 μM erastin (Supplemental Fig. S5). Erastin treatment strongly induced HR cell death, ROS and Fe^3+^ accumulation, and lipid peroxidation in *ΔOsmek2* mutant leaf sheaths at 72 and 96 hpi with avirulent *M. oryzae* 007, similar to that observed in rice DJ (Fig. 8 and Supplemental Fig. S6). By contrast, *M. oryzae* 007 infection induced disease-related cell death but not Fe^3+^ accumulation in erastin-untreated leaf sheaths of the susceptible *ΔOsmek2* mutant at 72 and 96 hpi (Fig. 8, A, C and D). However, the chemiluminescent assay indicated that the high ROS levels observed in the *ΔOsmek2* mutant were similar to those observed in rice DJ and erastin-treated *ΔOsmek2* mutant at 72 and 96 hpi (Fig. 8B). ROS accumulation, MDA peroxidation, and cell death phenotype were distinctly enhanced in the *ΔOsmek2* mutant; however, increased Fe^3+^ accumulation was not observed at 72 and 96 hpi (Fig. 8, A and C; Supplemental Fig. S6). Avirulent *M. oryzae* 007 infection did not increase the number of Prussian blue-stained cells in *ΔOsmek2* leaf sheaths at 72 and 96 hpi (Fig. 8D). These combined results indicate that disease-related cell death is ROS-dependent but iron-independent in the compatible rice–*M. oryzae* interaction. Increased ROS production and lipid peroxidation in *M. oryzae*-infected tissues may induce susceptibility-related cell death that facilitates subsequent fungal invasion and infection. However, intracellular iron accumulation may not be required for disease-related cell death in compatible rice–*M. oryzae* interactions.

**Figure 8.**
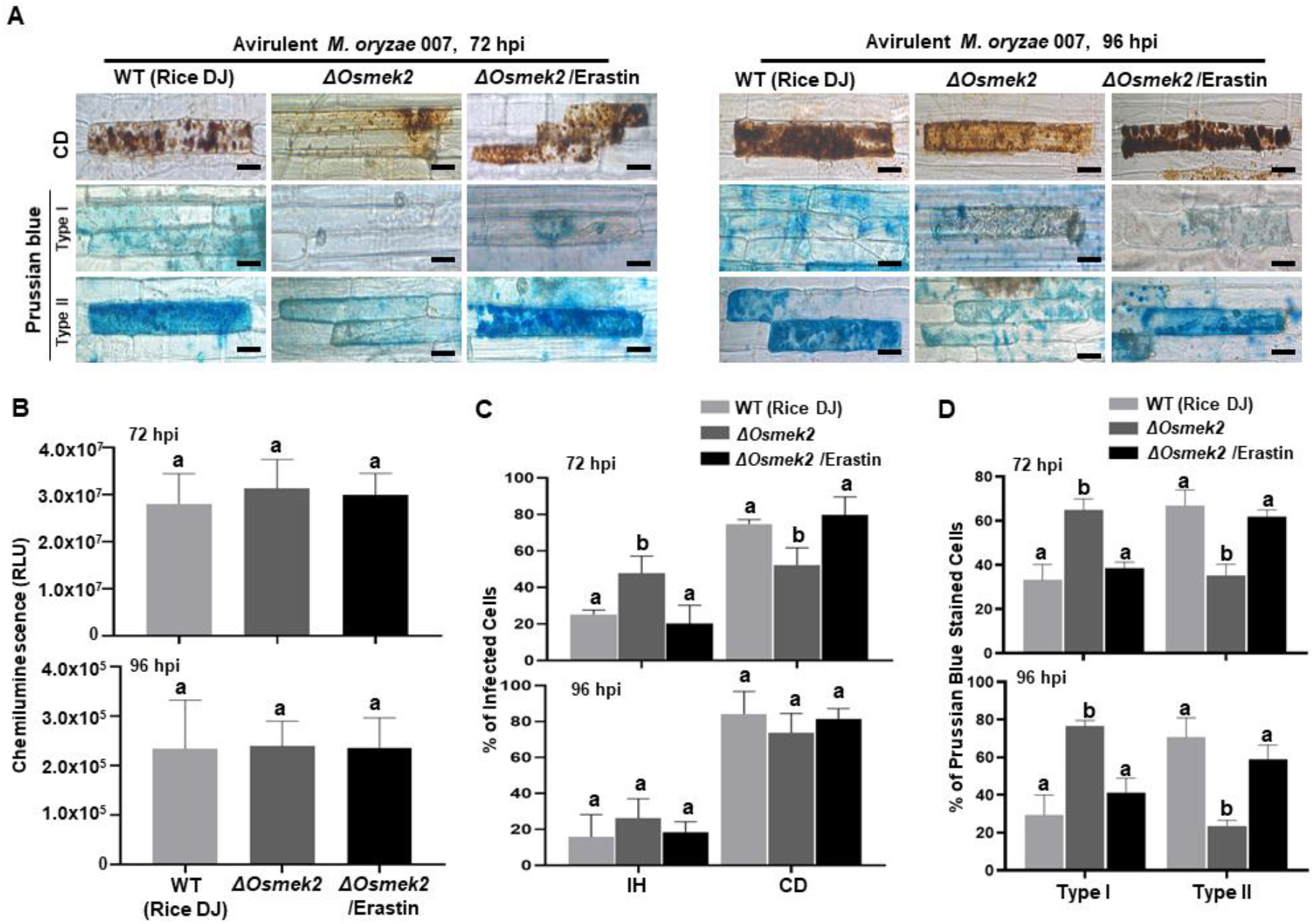
Disease-related cell death is ROS-dependent and iron-independent in the compatible *ΔOsmek2* mutant–*Magnaporthe oryzae* interaction. *ΔOsmek2* mutant leaf sheaths were inoculated with conidial suspensions (4×10^5^ conidia mL^−1^) of *M. oryzae* 007 containing 10 μM erastin. *M. oryzae* 007 infection induced disease-related cell death, but *ΔOsmek2* mutant leaf sheath cells did not accumulate ferric ions (Fe^3+^) at 96 h after inoculation. A, Erastin treatment induces cell death and ferric ion (Fe^3+^) accumulation in *ΔOsmek2* mutant leaf sheath cells during avirulent *M. oryzae* 007 infection. Prussian blue (blue color) staining shows ferric ion accumulation in rice cells. The images are representative of different leaf sheath samples from three independent experiments. Scale bars=20 μm. B, Quantification of ROS accumulation in leaf sheath cells. ROS quantities were monitored using a GloMax^®^ 96 Microplate Luminometer (Promega). Values are means ± SD of total relative luminescent units (RLU) (*n*=10). C, Quantification of infected cell phenotypes in rice leaf sheaths. Results are presented as mean values ± SD; *n*=4 leaf sheaths from different plants. D, Quantification of Prussian blue-stained cells. Prussian blue-stained cells were categorized into two phenotypes: Type I, cells that contain invasive hyphae (IH) but are weakly or not Prussian blue-stained; and Type II, strongly Prussian blue-stained with only a few poor hyphae. Images were taken using a fluorescence microscope (Zeiss equipped with Axioplan 2). Experiments were repeated three times with similar results. Results are presented as means ± SD; *n*=4 leaf sheaths. Different letters above the bars indicate significantly different means (*P*<0.05) as analyzed by Fisher’s protected LSD test. IH, invasive hyphae; CD, cell death; BF, bright field; GF, green fluorescence; hpi, hours post-inoculation.

### *OsMPK1* Overexpression Induces Iron- and ROS-Dependent Ferroptotic Cell Death in Rice during *M. oryzae* Infection

Rice MAP kinase (OsMPK1) is an interactor of OsMEK2 and actively involved in *M. oryzae* infection (Singh et al., 2012; Ueno et al., 2015). To investigate the involvement of OsMPK1 in ferroptotic cell death in rice, we overexpressed *OsMPK1* in the susceptible rice cultivar Nipponbarre (NB) under the control of CaMV 35S promoter. *OsMPK1* expression in rice NB and 35:*OsMPK1* mutant plants infected by virulent *M. oryzae* PO6-6 were analyzed using qRT-PCR (Supplemental Fig. S7). Filamentous and bulbous invasive hyphae (IH) remained viable and propagated to the neighboring cells in rice NB leaf sheaths. However, 35S:*OsMPK1* overexpression induced a hypersensitive cell death with poorly grown IH in leaf epidermal cells at 48 hpi (Fig. 9, A and B). DAB and CM-H_2_DCFDA staining showed accumulation of ROS around the IH and entire cell in 35S:*OsMPK1*; however, no or weak accumulation of ROS occurred in rice NB at 36-48 hpi during virulent *M. oryzae* PO6-6 infection (Fig. 9, A and D). Chemiluminescence assay with a luminometer revealed that ROS levels distinctly increased in 35S:*OsMPK1* mutant cells compared to rice NB cells at 48 hpi (Fig. 9E). Prussian blue staining of Fe^3+^ showed strong accumulation of ferric ion in 35S:*OsMPK1* mutant at 48 hpi (Fig. 9, A and C). We further analyzed lipid peroxidation in rice NB and 35S:*OsMPK1* mutant cells by detecting MDA production at 48 hpi (Fig. 9F). Lipid peroxidation levels were significantly higher in 35S:*OsMPK1* mutant cells than in rice NB cells. The combined results indicate that *OsMPK1* is involved in ROS and Fe^3+^ accumulation and lipid peroxidation leading to the ferroptotic cell death during *M. oryzae* infection.

**Figure 9.**
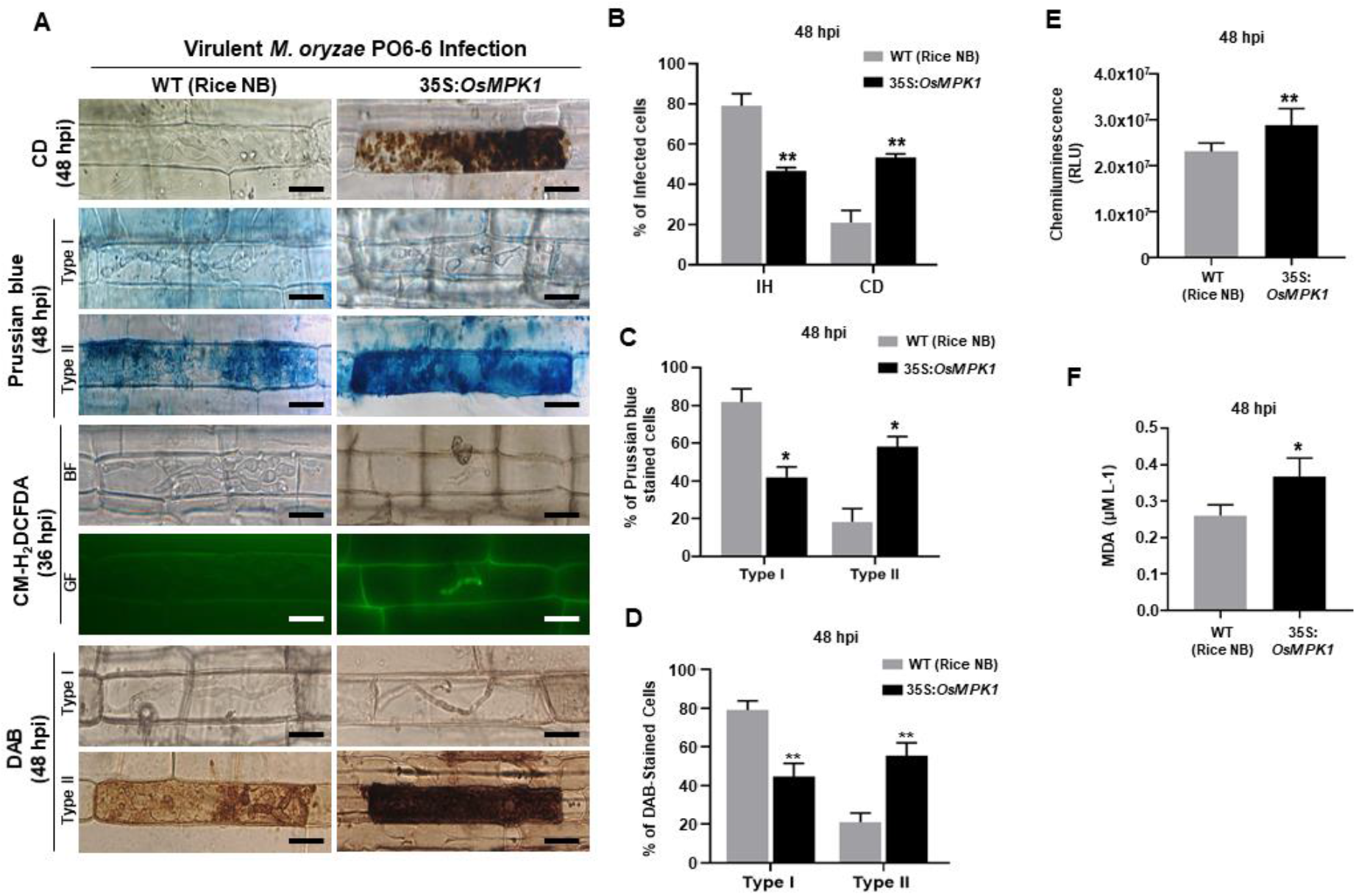
*OsMPK1* overexpression induces ROS and ferric Ion (Fe^3+^) accumulation, lipid peroxidation and cell death in rice leaf sheaths during virulent *Magnaporthe oryzae* PO6-6 infection. Rice leaf sheaths of the susceptible wild-type (WT) cultivar NB and *OsMPK1*-overexpressed (35S:*OsMPK1*) mutant plants were inoculated with the conidial suspension (4×10^5^ conidia/mL) of virulent *M. oryzae* PO6-6. A, Microscopic images of cell death and ROS and ferric Ion (Fe^3+^) accumulation in rice sheath cells at 48 hpi. ROS accumulation in the infected leaf sheath epidermal cells was detected by CM-H_2_DCFDA (green fluorescence) and DAB (dark brown color) staining. Prussian blue (blue color) staining shows ferric ion accumulation in rice cells. The images are representatives of different leaf sheath samples from three independent experiments. Scale bars=20 μm. B, Quantification of cell death (CD) and invasive hyphae (IH) in rice sheath cells at 48 hpi. C, Quantification of Prussian blue-stained cells. Prussian blue-stained cells were divided into two phenotypes: Type I, cells that contain invasive hyphae (IH) but are weakly or not Prussian blue-stained; and Type II, strongly Prussian blue-stained with only a few poor hyphae. D, DAB-stained cell phenotypes. DAB-stained cells were divided into two phenotypes: Type I, cells that contain invasive hyphae (IH) but are weakly or not DAB-stained; and Type II, strongly DAB-stained cells with only a few poor hyphae. E, Quantification of ROS accumulation. ROS accumulation was monitored using a GloMax^®^ 96 Microplate Luminometer (Promega). Values are means ± SD of total relative luminescent units (RLU) (*n*=10). F, Determination of lipid peroxidation by MDA (malondialdehyde) assay. Images were taken using a fluorescence microscope (Zeiss equipped with Axioplan 2) with bright field and green fluorescence (GF) filters. Experiments were repeated three times with similar results. Results are presented as mean values ± SD; *n*=4 leaf sheaths from different plants. Asterisks indicate statistically significant differences (Student’s *t*-test, *P<* 0.01). hpi, hours post-inoculation.

### *OsMEK2* Expression Positively Regulates *OsNADP-ME* and *OsRbohB* Expression in Rice during *M. oryzae* 007 Infection

We recently reported that rice NADP-malic enzyme (OsNADP-ME) and respiratory burst oxidase homolog (OsRboh, NADPH-oxidase) are involved in Fe^3+^ and ROS accumulation during cell death and defense responses in rice (Dangol et al. 2019). Interaction of *N. benthamiana* WRKY8 with MAPKs induce the downstream target genes *NADP-ME* and *Rboh*, resulting in the ROS burst (Yoshioka et al., 2003; Ishihama et al., 2011). RbohB activation via MAPK cascades is required for the pathogen-responsive ROS burst (Adachi and Yoshioka, 2015). Here, we analyzed the expression of *OsNADP-ME2-3* (Singh et al., 2016) and *OsRbohB* (Wong et al., 2007) in rice DJ, *ΔOsmek2* and 35S:*OsMEK2* mutant plants during avirulent *M. oryzae* infection (Fig. 10). *OsNADP-ME2-3* expression patterns did not differ in rice DJ and *ΔOsmek2*, except for a reduction in *ΔOsmek2* at 12 hpi. However, *OsNADP-ME2-3* was distinctly expressed in 35S:*OsMEK2* mutant plants at 12 and 72 hpi. *OsRbohB* expression was significantly downregulated in *ΔOsmek2,* but distinctly upregulated at 24 hpi, compared to that in rice DJ (Fig. 10). These combined results indicate that *OsMEK2* expression positively regulates *OsNADP-ME* and *OsRbohB* expression during avirulent *M. oryzae* infection.

**Figure 10.**
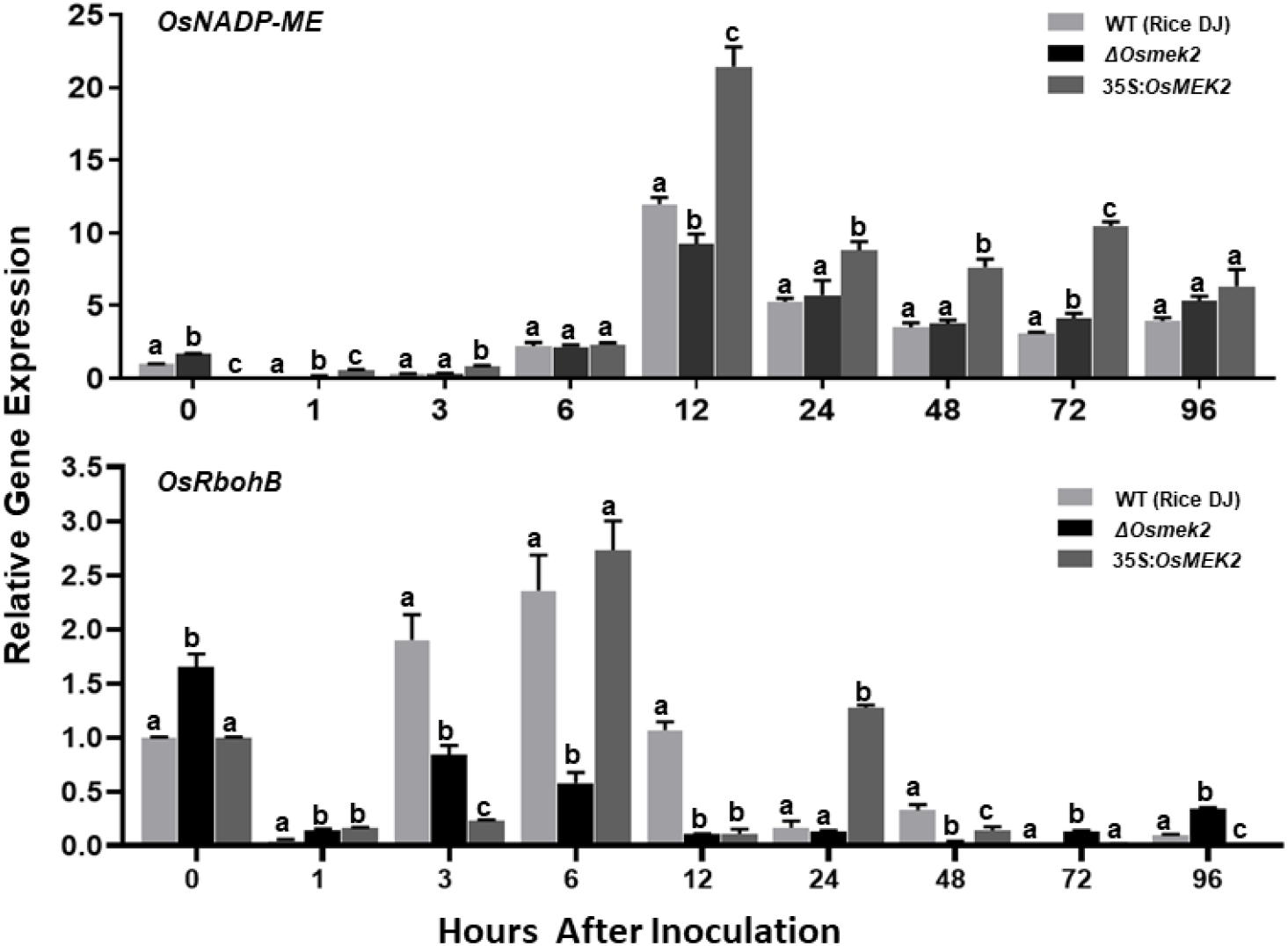
Quantitative real-time RT-PCR analysis of time-course expression of *OsNADP-ME* and *OsRbohB* in rice leaf sheaths infected with avirulent *Magnaporthe oryzae* 007. Leaf sheaths of the wild-type (WT) rice cultivar DJ, *OsMEK2-*deleted (*ΔOsmek2*) and *OsMEK2-*overexpressed (35:*OsMEK2).*mutant plants were sampled at different time points after inoculation, followed by total RNA extraction. Relative gene expression levels of *OsNADP-ME* and *OsRbohB* at each time point were calculated by normalizing with respect to the expression of the internal control *18S rRNA* gene. Data represent the means ± SD from three independent experiments. Different letters above the bars indicate significantly different means (*P*<0.05) as analyzed by Fisher’s protected LSD test.

### Subcellular Localization of OsMEK2, OsMPK1, and OsWRKY90

The subcellular localization study of MAP kinase signaling proteins is important for understanding their biological functions in plant cells. In this study, we investigated subcellular localization of green fluorescent protein (GFP)-tagged 35S:OsMEK2 (OsMEK2:GFP), OsMPK1:GFP, and OsWRKY90:GFP in *N. benthamiana* leaves using *A. tumefaciens*-mediated transient expression (Fig. 11). The nuclei inside cells were counterstained with DAPI to help verify nuclear localization of GFP-tagged proteins. The control GFP construct (00:GFP) was ubiquitously detected in the cytoplasm of *N. benthamiana* cells. OsMEK2:GFP was localized mainly to the cytoplasm, but also to some nuclei in *N. benthamiana* cells. OsMPK1:GFP was localized to both the cytoplasm and nuclei. However, the OsWRKY90:GFP transcription factor was located inside the nuclei, but not in the cytoplasm. These results indicate that OsMEK2 interacts with OsMPK1 in the cytoplasm, and OsMPK1 moves into the nuclei to interact with the OsWRK90 transcription factor.

**Figure 11.**
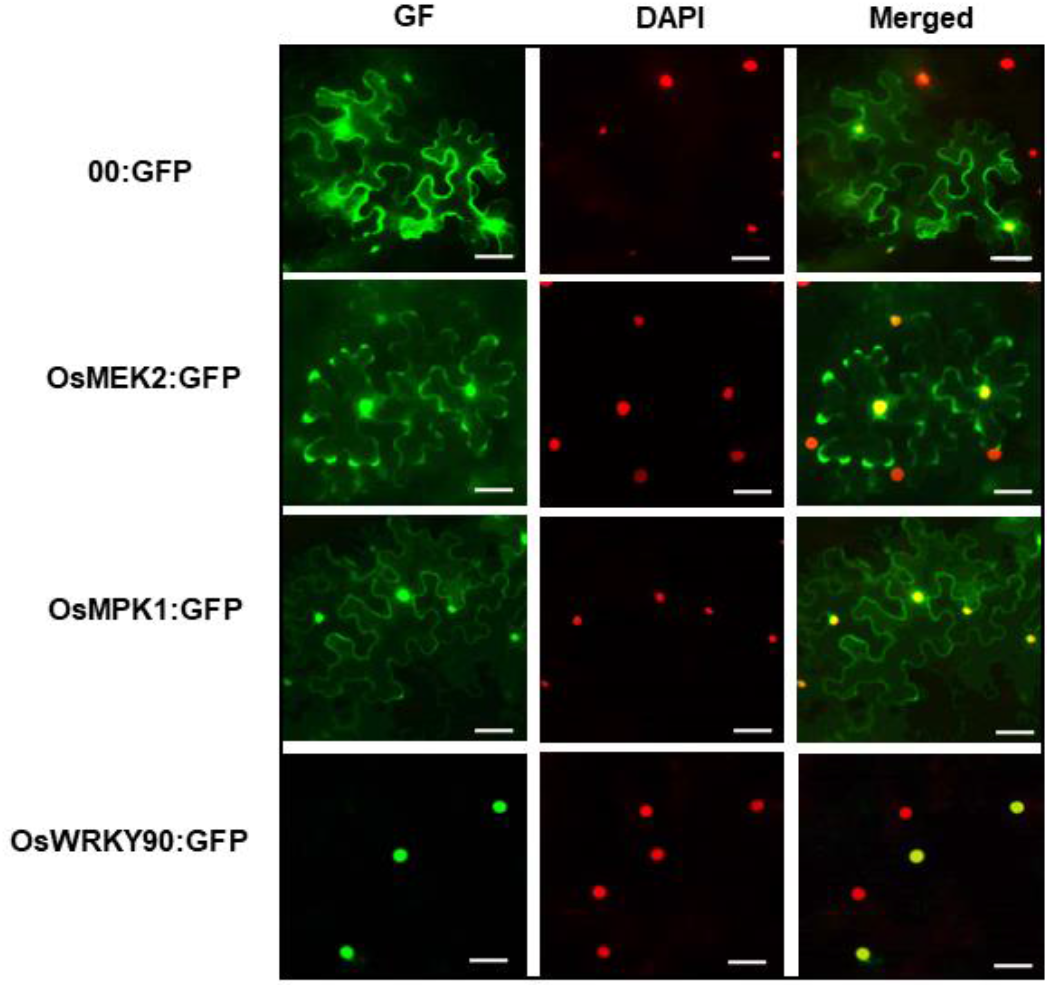
Subcellular localization of OsMEK2, OsMPK1, and OsWRKY90 at 36 h after agroinfiltration into *Nicotiana benthamiana* leaves. 4’,6-diamidino-2-phenylindole (DAPI) staining was used to visualize nuclei in *N. benthamiana* epidermal cells. Images of subcellular localization of 00:GFP, OsMEK2:GFP, OsMPK1:GFP, and OsWRKY90:GFP were taken with a fluorescence microscope using bright field, GF (green fluorescence) and DAPI filters. GFP, green fluorescent protein. Scale bars *=*50μm.

## DISCUSSION

Plant mitogen-activated protein kinase (MAPK) cascades are involved in signaling multiple defense responses, the HR, and cell death responses during pathogen invasion and infection (Meng and Zhang, 2013; Thulasi Devendrakumar et al., 2018). We recently reported a ferroptotic cell death response in rice (*Oryza sativa*) during *Magnaporthe oryzae* infection (Dangol et al., 2019). Ferroptosis is a form of nonapoptotic iron-dependent cell death that was first discovered in oncogenic mammalian cells (Dixon et al., 2012). OsMEK2 interacts with OsMPK1 (Singh et al., 2012). Here, we demonstrated that rice MAP kinase (OsMEK2 and OsMPK1) signaling was required for iron- and ROS-dependent ferroptotic cell death in rice–*M. oryzae* interactions, and blast disease (susceptibility)-related cell death was ROS-dependent but iron-independent in the susceptible *ΔOsmek2* mutant plants.

### Rice MAPK Kinase 2 (OsMEK2) is Involved in the MAPK Signaling Network Leading to HR-Mediated Resistance to *M. oryzae* Infection

We previously reported that OsMEK2 physically interacts with and phosphorylates downstream OsMPK1 and OsMPK6 (Singh et al. 2012). MAPK kinase (MEK)-MAPK interactions may have functional roles in HR cell death responses and MAPK signaling networks during *M. oryzae* infection in rice plants. OsMEK2 shares relatively high sequence homology with both AtMKK1 and AtMKK2. Arabidopsis MAPK cascades are involved in innate immunity responses (Rasmussen et al., 2012). Arabidopsis MAPKKs phosphorylate downstream MAPKs and WRKY transcription factors. OsMAPKK10-2 physically interacts with and phosphorylates OsMAPK6 and OsMPK3 (Ma et al., 2017).

In the present study, avirulent *M. oryzae* 007 infection triggered *OsMEK2* expression in the resistant DJ WT rice cultivar at early time points after inoculation. *OsMEK2* deletion in DJ plants induced a susceptible (disease) response to avirulent *M. oryzae* infection in *ΔOsmek2* mutant plants; however, *OsMEK2* overexpression in 35S:*OsMEK2* mutant plants redeemed hypersensitive cell death response against avirulent *M. oryzae* 007 infection. These combined results suggest that HR-mediated resistance signaling of OsMEK2 may occur during the incompatible rice–*M. oryzae* interaction. *OsMEK2* deletion and overexpression differentially regulated *OsMPK1*, *OsMPK6*, and *OsWRKY90* expression in *ΔOsmek2* and 35S:*OsMEK2* mutant plants, especially during early stages of *M. oryzae* infection. These results suggest that *OsMEK2* expression distinctly induced the downstream *OsMPK1* and *OsMPK6* signaling responses to avirulent *M. oryzae* infection. The pathogen-responsive MAPKs, OsMPK1 and OsMPK6 may be involved in hormonal signaling in disease resistance networks (Singh and Jwa, 2013; Ueno et al., 2015). We previously reported that OsMPK1 physically interacts with the OsWRKY80 (Singh et al., 2012) and OsWRKY90 (Shen et al., 2012) transcription factors. In plant disease resistance networks, WRKY transcription factors can associate with MAP kinases in the nuclei and regulate downstream defense-related gene expression (Pandey and Somssich, 2009; Ishihama et al., 2011; Jalmi and Sinha, 2016). *OsMEK2* expression distinctly induced pathogenesis-related protein 1b (*OsPR-1b*) but not ascorbate peroxidase1/2 (*OsAPX1/2*) during avirulent *M. oryzae* infection in rice. PR-1 proteins are markers of defense responses to pathogen infection in rice (Mitsuhara et al., 2008). Thus, *OsMEK2* signaling may trigger the MAP kinase cascade pathways leading to HR-mediated resistance to avirulent *M. oryzae* infection in rice.

### OsMEK2 and OsMPK1 Signaling Cascades Are Required for Iron- and ROS-Dependent Ferroptotic Cell Death Response

MAP kinase signaling cascades are highly conserved in diverse plant species and involved in plant defense responses (Tanaka et al., 2009; Melech-Bonfil and Sessa 2011; Oh et al., 2013; Ma et al., 2017). Activation of the MAPKK-MAPK cascades is associated with programmed cell death (PCD) in plants (Thulasi Devendrakumar et al., 2018). ROS mediate cellular defense responses against pathogen invasion in plants (Apel and Hirt, 2004; Mittler et al., 2004). Ferric ion (Fe^3+^) is essential in plants for HR cell death and defense- and disease-related iron homeostasis (Liu et al., 2007; Dangol et al., 2019). Iron is required for intracellular lipid peroxide accumulation (Stockwell et al., 2017; Dangol et al., 2019). Iron- and ROS-dependent ferroptotic cell death was first discovered in rice cells infected with different avirulent *M. oryzae* strains (Dangol et al., 2019).

Pathogen-responsive MAPKs may trigger the early ROS burst during plant defense and cell death responses (Meng and Zhang, 2013). Iron and ROS accumulation was not induced in *ΔOsmek2* mutant leaf sheaths, compared to WT DJ and 35S:*OsMEK2* mutant leaf sheaths during early *M. oryzae* infection (30–48 hpi). These results suggest that *OsMEK2* activation is one of the earliest signaling events involved in iron- and ROS-dependent ferroptotic cell death in rice. The ROS burst in rice may originate from the plasma membrane NADPH-oxidase (*OsRbohB*), which is activated during early *M. oryzae* infection (3–12 hpi). MAPKs could phosphorylate WRKY transcription factors to subsequently activate NADPH oxidases (Rbohs), which are essential for potent and prolonged ROS burst (Adachi et al., 2015; Jwa and Hwang, 2017). Early MAPK kinase (*OsMEK2*) signaling seems likely to activate *OsMPK1*, *OsWRKY90*, and NADPH-oxidase (*OsRbohB*), ultimately leading to the iron- and ROS-dependent ferroptotic cell death response.

In our study, overexpression of 35S:*OsMPK1* in susceptible rice cultivar NB significantly induced iron and ROS accumulation at 48 h after virulent *M. oryzae* PO6-6 infection. Ma et al., (2017) demonstrated that overexpression of rice *MPK6* (*OsMPK1* in current study) reduced susceptibility in rice cultivar Zhonghua 11 against *Xanthomonas oryzae pv. oryzae* infection. Our combined results suggest that the OsMEK2-OsMPK1-OsWRKY90 cascades positively regulate ferroptotic cell death in rice against *M. oryzae* infection. MEK2–SIPK/WIPK cascades triggered Pto/Prf-mediated ETI in both tomato and *Nicotiana benthamiana* (Melech-Bonfil and Sessa, 2011). Rice MPKK10.2 and MPK6 cascades induced resistance against *X. oryzae pv. oryzae* infection (Ma et al., 2017). Plant MAP kinases have been demonstrated to differentially regulate WRKY transcription factors in defense-related signaling pathways (Eulgem and Somssich, 2007, Pandey and Somssich, 2009; Ishihama and Yoshioka, 2012). Jalmi and Shina (2016) reported positive involvement of the OsMKK3-OsMPK7-OsWRKY30 module in inducing rice resistance against *X. oryzae* infection. Rice OsMPK7 interacts with and phosphorylates OsWRKY30 to mediate resistance against *X. oryzae* infection (Jalmi and Sinha, 2016).

Iron and ROS accumulation is required for lipid peroxidation to trigger ferroptotic HR cell death during avirulent *M. oryzae* infection in rice (Dangol et al., 2019). We observed that lipid peroxidation was significantly lower in the *ΔOsmek2* mutant than that in rice DJ and 35S:*OsMEK2* mutant plants. 35S:*OsMPK1* overexpression in susceptible rice NB plants triggered a high level of lipid peroxidation in rice leaf sheath cells. Theses combined results suggest that *OsMEK2-OsMPK1* signaling induces the accumulation of lipid-based ROS such as lipid peroxides. Lipid peroxide accumulation is iron-dependent, and ultimately leads to ferroptotic cell death (Stockwell et al., 2017). In mammals, intracellular iron reacts with lipid ROS inside oncogenic cells to trigger ferroptotic cell death (Dixon et al., 2012). In wheat (*Triticum aestivum*), iron deposition at the pathogen infection site mediates the ROS burst inside epidermal cells infected with powdery mildew (Liu et al., 2007).

### Erastin Triggers Iron- and ROS-Dependent Ferroptotic Cell Death in *ΔOsmek2* Mutant Rice Plants

Erastin is an oncogenic RAS-selective lethal (RSL) small molecule that effectively damages human cancer cells but does not affect isogenic normal cells (Dolma et al., 2003). Dixon et al. (2012) first discovered that erastin induces cellular iron-dependent lipid ROS accumulation in mammalian cells, leading to the unique iron-dependent nonapoptotic cell death (ferroptosis) in carcinogenic RAS-mutated cells. The ferroptosis inducer erastin inhibits glutathione peroxidase 4 (GPX4) activity to elevate cytoplasmic lipid ROS levels (Yang and Stockwell, 2016; Stockwell et al., 2017). GPX4 is an inhibitor of lipid peroxidation (Ursini et al., 1982), and reduces membrane phospholipid hydroperoxides to suppress ferroptosis (Stockwell et al., 2017).

In this study, we showed that erastin treatment of *ΔOsmek2* mutant rice triggered iron (Fe^3+^) and ROS accumulation and lipid peroxidation, leading to iron- and lipid ROS-dependent ferroptotic cell death during *M. oryzae* infection. These results suggest that erastin-induced ferroptotic cell death in rice is iron- and lipid ROS-dependent, but is independent of the rice MAPK kinase OsMEK2. In our earlier study, we validated these results by showing that erastin treatment triggered *OsMADP-ME2*– independent ferroptotic cell death in rice during *M. oryzae* infection (Dangol et al. 2019). NADP-ME provides the cytoplasmic electron donor NADPH for ROS production (Singh et al. 2016; Jwa and Hwang, 2017). Erastin treatment induces iron- and ROS-dependent ferroptotic HR cell death in compatible rice–*M. oryzae* interactions (Dangol et al., 2019). Thus, erastin-mediated induction of ferroptotic cell death in rice may not require specific cell death-related plant genes, such as *OsMEK2* and *OsNADP-ME2*. Plant and mammalian genes that are specifically regulated by erastin to trigger ferroptotic cell death have not yet been identified (Stockwell et al. 2017; Hirschhorn and Stockwell, 2019). Erastin-induced ferroptotic cell death in rice does not appear to be genetically controlled, but may occur nonspecifically. Future studies will investigate whether distinct genetic networks govern erastin-induced ferroptotic cell death in plants.

### Disease-Related Cell Death Is ROS-Dependent and Iron-Independent in *ΔOsmek2* Mutant Rice Plants

Disease-related cell death occurs during compatible (susceptible) interactions between plants and pathogens (Greenberg, 1997; Richberg et al., 1998). In this study, we showed that *M. oryzae* infection induced disease-related cell death that was not dependent on iron accumulation in susceptible rice cells at the late infection stage, when typical disease symptoms are visible in infected rice. ROS accumulation, lipid peroxidation, and cell death phenotypes distinctly increased in the *ΔOsmek2* mutant at the late *M. oryzae* 007 infection stages (72 and 96 hpi); however, significant iron accumulation did not occur. Iron-independent and ROS-dependent cell death at late infection stages in the compatible rice–*M. oryzae* interaction is distinct from ferroptotic HR cell death, but similar to necrosis-like cell death. Necrotic cell death caused by compatible plant interactions with necrotrophic pathogens is dependent on ROS accumulation (Mengiste, 2012).

The compatible interaction between rice and *M. oryzae* triggers significantly low ROS and iron accumulation leading to a low level of ferroptotic cell death, which subsequently facilitates *M. oryzae* hyphae to lively propagate into the neighboring cells (Dangol et al., 2019). However, compatible interactions of plants with pathogens cause disease symptoms, which are accompanied by disease-related cell death at late infection stages (Greenberg, 1997; Gilchrist, 1998). A toxin or secreted virulence factor from the microbial pathogen may directly kill plant cells or trigger an endogenous cell death program (Greenberg, 1997). Cell death in compatible interactions may derive from pathogen-mediated necrosis rather than host-induced PCD (Morel and Dangl, 1997; Gilchrist, 1998). ROS accumulation and lipid peroxidation during *M. oryzae* infection at the late stages (i.e., 96 hpi) are involved in disease (susceptibility)-related cell death in rice, as suggested previously (Govrin and Levine, 2000; Greenberg and Yao, 2004; Torres et al., 2006; Choi et al., 2013; Jwa and Hwang, 2017). However, intracellular iron accumulation may not be required for disease-related cell death in compatible rice–*M. oryzae* interactions. By contrast, iron accumulation is likely essential for the induction of ferroptotic cell death to restrict avirulent *M. oryzae* invasion into rice cells. How cell death in compatible interactions differs from HR cell death caused by incompatible interactions remains to be fully elucidated.

### Proposed Model of OsMEK2-OsMPK1-OsWRKY90 Signaling Pathways in Ferroptotic Cell Death during Rice*–M. oryzae* Interactions

We previously reported that avirulent *M. oryzae* infection induces iron- and ROS-dependent ferroptotic cell death in rice (Dangol et al., 2019). Here, we combine our cumulative data to propose the following working model: OsMEK2-OsMPK1-OsWRKY90 signaling positively regulates iron- and ROS-dependent ferroptotic HR cell death in rice–*M. oryzae* interactions (Figure 12). The invasion of avirulent *M. oryzae* 007 into rice cells activates rice MAP kinases (*OsMEK2* and *OsMPK1*) via different hypothesized MAPK signaling pathways.

**Figure 12.**
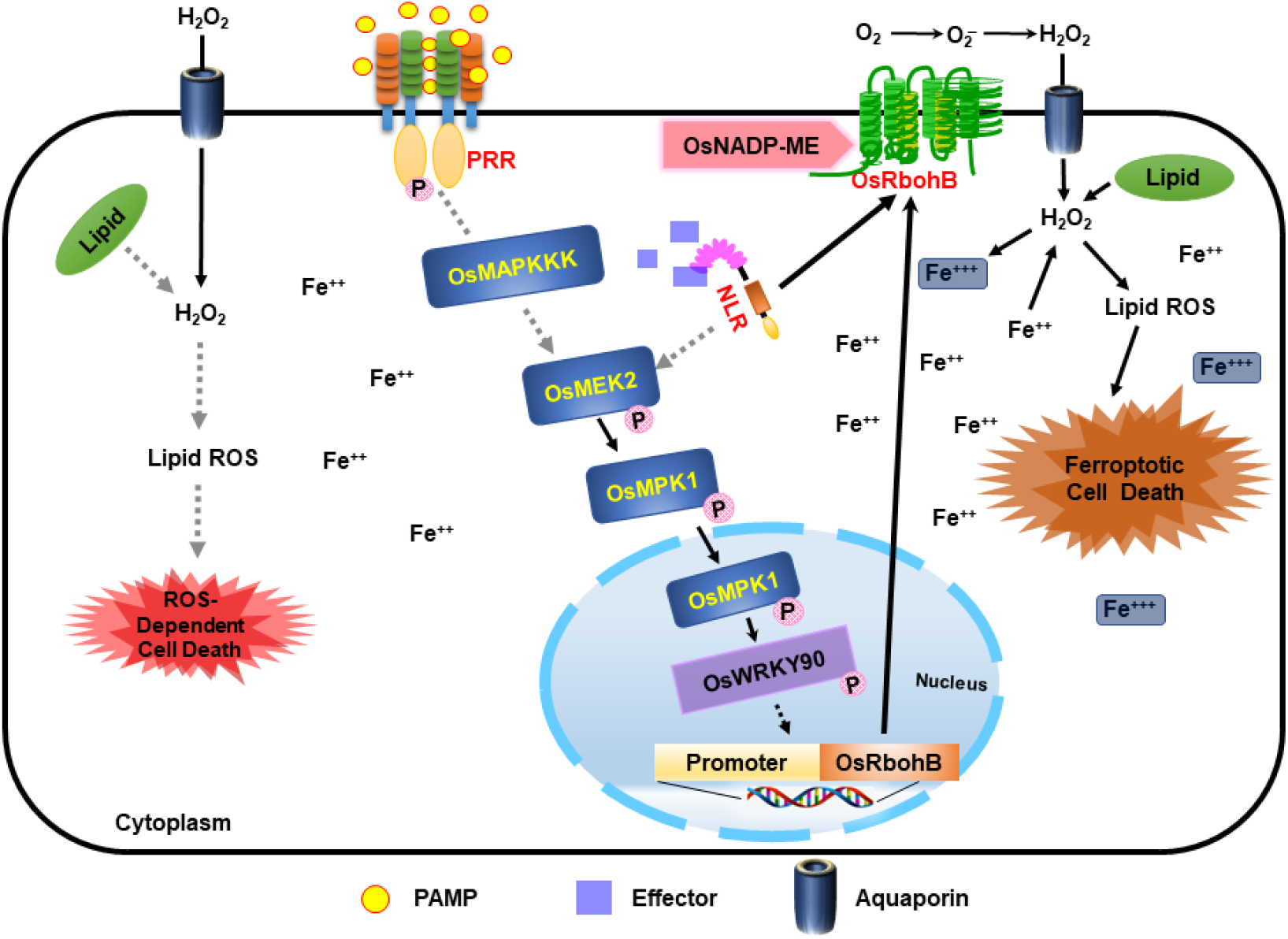
Proposed model of rice MAP kinase kinase 2 (OsMEK2) signaling pathways leading to iron- and ROS-dependent ferroptotic cell death in incompatible rice–*Magnaporthe oryzae* interactions. *M. oryzae* invasion in rice cells activates rice MAP kinase kinase 2 (OsMEK2) via different hypothesized signaling pathways (gray dotted lines). The perception of PAMPs or pathogen effectors via membrane-bound PRRs or NLRs, respectively, activates MAP kinases in plant cells. Active OsMEK2 triggers OsMPK1-OsWRKY90 pathways in the nucleus, which may lead to upregulation of NADP-malic enzyme (ME) and NADPH-oxidase (*OsRbohB*). The *de novo* synthesis of OsRbohB and its trafficking to the plasma membrane contributes to iron- and ROS-dependent ferroptotic death in rice cells. Disease (susceptibility)-related cell death is ROS-dependent and iron-independent in the compatible *ΔOsmek2* mutant–*M. oryzae* interaction. MEK, mitogen-activated protein kinase kinase; MPK, mitogen-activated protein kinase; NADP-ME, NADP-malic enzyme; NLR, nucleotide-binding leucine-rich repeat; PAMP, pathogen-associated molecular pattern; PRR, pattern recognition receptor; WRKY, tryptophan (W), arginine (R), lysine (K), tyrosine (Y) transcription factor.

The perception of PAMPs or pathogen effectors via membrane-bound PRRs or NLRs, respectively, activates OsMAP kinase cascades in rice cells, as proposed previously (Jones and Dangle, 2006; Zipfel, 2008). MAP kinase kinases (MEKs) activate MAP kinases, which migrate from the cytoplasm to the nucleus and regulate transcriptional reprogramming (Morris, 2001; Ahlfors et al., 2004). OsMEK2 interacts with and phosphorylates downstream rice MAP kinase 1 (OsMPK1) and MAP kinase 6 (OsMPK6) in the cytoplasm (Singh et al., 2012). OsMPK1 moves from the cytoplasm into the nucleus to interact with the OsWRKY90 transcription factor (Figure 11, Singh et al., 2012). *OsMEK2* expression triggers OsMPK1-OsWRKY90 signaling pathways in the nucleus (Figure 11), which may lead to the upregulation of OsNADP-malic enzyme and rice NADPH-oxidase B (*OsRbohB*) (Fig. 12). The MAPK-WRKY pathway activates Rbohs, leading to a prolonged and robust ROS burst (Adachi et al., 2015). The *de novo* synthesis of OsRbohB and its trafficking to the plasma membrane is involved in iron- and ROS-dependent ferroptotic death in rice cells. Rice MAP kinase 1 (OsMPK1) also may target OsWRKY90 to bind to specific sequences of some defense-related genes such as *OsPR-1b*.

Plant Rbohs have crucial roles in ROS (H_2_O_2_) production via cytosolic NADPHs supplying electrons to cell membranes (Suzuki et al., 2011; Kadota et al., 2015). The superoxide (O_2–_) produced from apoplastic oxygen (O_2_) can be converted to hydrogen peroxide (H_2_O_2_) by superoxide dismutase (SOD) (Fig. 12; Marino et al., 2012; Kadota et al., 2015). Apoplastic ROS (H_2_O_2_) produced by plasma membrane-bound Rbohs during ETI migrates across the plasma membrane using aquaporin channels and into the cell (Bienert and Chaumont, 2014; Jwa and Hwang, 2017). The simultaneous accumulation of H_2_O_2_ and Fe^2+^ in infected rice cells can cause a fenton reaction, in which ROS reacts with highly active Fe^2+^ to produce Fe^3+^ and highly toxic hydroxyl radicals (^●^OH) (Fenton, 1984; Pierre and Fontecave, 1999). The increased accumulation of iron (Fe^3+^) and lipid ROS triggers lipid peroxidation and subsequent ferroptotic cell death (Fig. 12; Dixon et al., 2012, Stockwell et al., 2017; Dangol et al., 2019).

Disease (susceptibility)-related cell death is lipid ROS-dependent, but iron-independent, in the compatible rice–*M. oryzae* interaction (Fig. 12). ROS accumulation, lipid (MDA) peroxidation, and cell death phenotype were significantly enhanced in susceptible *ΔOsmek2* mutant plants at later stages of *M. oryzae* 007 infection (72 and 96 hpi). However, iron (Fe^3+^) did not accumulate in the *ΔOsmek2* mutant during *M. oryzae* 007 infection. Iron accumulation may not mediate disease-related cell death in compatible rice–*M. oryzae* interactions. ROS accumulation was detected in plant cells during disease-related cell death at later stages of pathogen infection in susceptible plant–necrotrophic pathogen interactions (Mengiste, 2012).

This study provides evidence for the significance of OsMEK2-OsMPK1-OsWRKY90 signaling cascades during the induction of ferroptotic cell death in rice– *M. oryzae* interactions. Rice *OsMEK2* expression positively regulates iron and ROS accumulation in avirulent *M. oryzae*-infected cells. Iron- and ROS-dependent ferroptotic cell death in rice is a generally regulated form of cell death that is common in incompatible rice–*M. oryzae* interactions, irrespective of rice and *M. oryzae* genotypes (Dangol et al., 2019). Iron accumulation in ferroptotic cells may be harmful to the invaded hyphae of avirulent *M. oryzae*. Iron-independent disease-related cell death in compatible rice–*M. oryzae* interactions is likely a necrosis-type cell death that is distinct from ferroptotic cell death in incompatible rice–*M. oryzae* interactions. However, further research on the functions of this ROS-dependent disease-related cell death is required to determine how virulent *M. oryzae* infection suppresses ferroptotic cell death and induces disease-related cell death, which may be beneficial for pathogen growth *in planta*.

## MATERIALS AND METHODS

### Plant Material and Growth Conditions

The WT rice cultivars Dongjin (DJ) and Nipponbarre (NB) the *ΔOsmek2* mutant plant were used in this study. *ΔOsmek2* T-DNA insertion mutant seeds were provided by the Rice Functional Genomic Express Database (RiceGE) managed by the Salk Institute Genomic Analysis Laboratory (http://signal.salk.edu./cgi-bin/RiceGE) (Jeon et al., 2000). DJ rice seeds were obtained from the National Institute of Crop Science, Korea (http://www.nics.go.kr). Rice seeds were germinated in water for 5 days and then planted in plastic pots containing Baroker soil (Seoul Bio, Korea). Rice plants were grown in growth chambers at 28°C under white fluorescent light (150 μmol photons m^−2^ s^−1^) with a 16 h photoperiod and 60% relative humidity.

### Fungal Cultures and Growth Conditions

The rice blast fungal strains *Magnaporthe oryzae* 007 and PO6-6 were provided by the Center for Fungal Genetic Resources, Seoul National University, Seoul, Korea (http://genebank.snu.ac.kr). *M*. *oryzae* 007 was avirulent (incompatible) and *M*. *oryzae* PO6-6 was virulent (compatible) to the rice cultivar DJ. The fungal cultures were stored at –20°C and cultured on rice bran agar media (20 g rice bran, 20 g sucrose, and 20 g agar in 1 L Milli-Q water). *M. oryzae* strains were grown at 25°C in the dark for 2 weeks. *M. oryzae* sporulation was induced by removing aerial mycelia from the fungal culture plates, followed by their incubation under a continuous fluorescent light (80 μmol photons m^−2^ s^−1^) for 2–3 days at 25°C.

### Fungal Inoculation of Rice Tissues and Infection Evaluation

Conidial suspensions of *M. oryzae* strains were inoculated on rice leaves and leaf sheaths as described previously (Singh et al., 2016; Dangol et al., 2019). *M. oryzae* conidia were harvested from the sporulated culture plates using a 0.025% Tween 20 (Sigma-Aldrich) solution. The conidial concentration was adjusted to 4×10^5^ conidia mL^−1^. The conidial suspension was spray-inoculated over the surface of 2-week-old rice seedlings. The inoculated seedlings were incubated at 25–28°C for 24 h under dark and moist conditions, and then were moved to normal conditions (16 h light/8 h dark). Disease phenotypes were observed at 5 days after inoculation and classified with respect to susceptible (large, elliptical, grayish, and expanded lesions) and resistant (slightly elongated, necrotic brownish spots) reactions.

Middle-aged leaf sheaths (5–7 cm lengths) of 4- or 5-week-old rice plants were inoculated with *M*. *oryzae* conidial suspensions (4 × 10^5^ conidia mL^−1^). Inoculated leaf sheaths were incubated in a moistened box with 100% relative humidity at 25°C under dark conditions. After incubation for different times, epidermal layers were excised from the leaf sheaths, cut into 1.5 cm lengths, and fixed on glass microscope slides. The infected epidermal cells were observed under the microscope and divided into two infection phenotypes: cells with viable IH and cells with HR cell death. The number of cells with each phenotype were counted at least three times from one representative of three independent experiments.

### *Identification of T-DNA Insertion in* ΔOsmek2 *Mutants*

*ΔOsmek2* T-DNA insertion mutant seeds from RiceGE (Jeon et al., 2000) were screened by PCR using the left gene-specific primer (LP), the right gene-specific primer (RP), and the T-DNA right border primer (RB). The gene-specific primers are listed in Supplemental Table S1. To verify homozygosity in the mutant plants, PCR was performed with LP and RP primers of the gene, and homozygous plants were identified by the lack of specific PCR products. The LP and RB primers were used for PCR analysis to confirm the presence of the T-DNA insertion in *ΔOsmek2* mutant plants. RT-PCR and immunoblotting analyses were performed to verify transcriptional and translational expression of *OsMEK2* in *ΔOsmek2* mutants.

### Rice Transformation

The full length cDNAs of *OsMEK2* and *OsMPK1* were amplified from rice cDNA library and inserted into plant expression vector pCAMLA under the control of CaMV 35S promoter, followed by selection of hygromycin gene (*hph*). The constructed CaMV 35S:*OsMEK2* and CaMV 35S:*OsMPK1* were introduced in the rice cultivars DJ and NB, respectively, by *Agrobacterium tumenfaciens-*mediated transformation, as described previously with slight modification (Hiei et al., 1994).

Briefly, 35S:*OsMEK2* and 35S:*OsMPK1* were delivered into rice calli using *A. tumenfaciens* strain LBA4404 (Hoekema et al., 1983; Lee et al., 2005). The transformed calli were selected on the selection media containing gradually increasing concentrations of hygromycin (30 mg/L and 60 mg/L). After rooting and shooting, rice seedlings were transferred in water for 4 days. After adaption in water, rice seedlings were raised in soil in a growth chamber. The positive transformants from T_0_ generation were selected by PCR using hygromycin primers. Next, seeds of T_1_ generation were analyzed on the hygromycin-containing media and T_2_ generation seeds were used for homozygote selection. The functional analysis was performed from T_3_ generation. The levels of gene expression were determined by immunoblot analysis and qRT-PCR. The primers used for the experiments are listed in Supplemental Table S1.

### Real-Time RT-PCR Analyses

Total RNA was isolated from rice tissue using TRIzol reagent (Invitrogen) according to the manufacturer’s instructions. The gene expression levels were analyzed by reverse-transcription PCR (RT-PCR) and real-time quantitative PCR (qRT-PCR). First-strand cDNA was synthesized from 2 μg total RNA in 20 μL reaction mixture using a cDNA synthesis kit (Invitrogen) according to the manufacturer’s instructions. Prepared cDNA (1 μL) was used as a template for both RT-PCR and qRT-PCR. The qRT-PCR was performed using TOPreal™ qPCR 2× PreMIX (SYBR Green with low ROX; Enzynomics, Daejeon, South Korea) according to the manufacturer’s instructions. Relative gene expression levels were determined using rice 18S ribosomal RNA or rice ubiquitin as an internal standard gene. Gene-specific primers used for the real-time RT-PCR analysis are listed in Supplemental Table S1.

### Protein Isolation and Immunoblot Assay

Rice proteins were extracted using trichloroacetic acid (TCA)/acetone extraction buffer [TCAAEB; 10% (w/v) trichloroacetic acid and 0.07% ß-mercaptoethanol in 100 mL acetone] as described previously (Cho et al., 2006). Rice leaves were ground to a fine powder using liquid nitrogen in a mortar and pestle. Then, proteins were precipitated with TCAAEB and washed three times with wash buffer [0.07% ß-mercaptoethanol, 2 mM ethylenediaminetetraacetic acid (EDTA), and EDTA-free protease inhibitor cocktail tablet (Roche) in a final volume of 100 mL acetone]. Protein precipitates were air-dried at room temperature, stored at –80°C for at least 24 h, and solubilized in lysis buffer containing thiourea and Tris (LB-TT) {7 M urea, 2 M thiourea, 4% (w/v) 3-[(3-cholamidopropyl) dimethylammonio]-l-propanesulfonate (CHAPS), 18 mM Tris-HCI (pH 8.0), 14 mM Trizma base, two EDTA-free protease inhibitor cocktail tablets (Roche), 0.2% (v/v) Triton X-100, and 50 mM dithiothreitol (DTT) in a final volume of 100 mL}. After centrifuging at 15,000×*g* for 15 min at 4°C, the supernatants were precipitated using pre-chilled acetone and solubilized in LB-TT buffer.

The OsMEK2 protein expression levels in DJ and *ΔOsmek2* mutant plants were determined by 10% SDS-polyacrylamide gel electrophoresis (PAGE) followed by immunoblot analysis using rabbit polyclonal anti-MEK2 antibody (EnoGene^®^ E580135-A-SE).

### Erastin Treatment

The small molecule cell death inducer erastin was used to investigate whether erastin treatment triggered ferroptotic cell death in *ΔOsmek2* mutant leaf sheath cells as described previously (Dangol et al., 2019). The harvested *M. oryzae* conidia (4×10^5^ conidia mL^−1^) were mixed with 10 μM erastin (Sigma-Aldrich, St. Louis, MO) and inoculated on leaf sheaths. The erastin-treated and *M. oryzae*-inoculated leaf sheath tissues were incubated in the dark at 25°C.

### CM-H2DCFDA Assay and DAB Staining

Cellular ROS (H_2_O_2_) localization in rice leaf sheath cells was visualized using 5-(and 6-) chloromethyl-2′,7′-dichlorofluorescin diacetate acetyl ester (CM-H_2_DCFDA) and 3,3′-diaminobenzidine (DAB) staining methods as described previously (Shin et al., 2005; Dangol et al., 2019). Briefly, thin epidermal layers of rice leaf sheaths were excised and cut into equal pieces, followed by incubation in 1 mL water for 5 min to remove wound-induced ROS. Epidermal sheath samples were incubated in 2 μM CM-H_2_DCFDA (Molecular Probes Life Technologies, Eugene, OH) in 1× phosphate-buffered saline (PBS) buffer in the dark for 30 min on a horizontal shaker. The incubated sheath samples were washed twice with 1× PBS buffer for 5 min in the dark. ROS localization inside the epidermal sheath cells was observed immediately under a fluorescence microscope.

For DAB staining, epidermal layers of rice leaf sheaths were vacuum-infiltrated with 1 mg mL^−1^ DAB (Sigma-Aldrich, St. Louis, MO) solution for 5 min, followed by overnight destaining with ethanol:acetic acid:glycerol (3:1:1, v/v/v). ROS localization in the DAB-stained epidermal cells was observed under a microscope. The DAB-stained cells were categorized into two phenotypes: Type I, infected cells that display no or weak DAB staining; and Type II, infected cells that display strong DAB staining.

### Chemiluminescence Assay for ROS Measurement

The chemiluminescence assay was used to measure ROS production in rice leaf sheaths as described previously (Singh et al., 2016; Dangol et al., 2019) with minor modifications. Epidermal layers of treated and *M. oryzae*-inoculated rice leaf sheaths were cut into small pieces (0.5 cm length) and incubated in 1 mL of sterilized Milli-Q water for 5 min to remove wound-induced ROS. Then, a piece of epidermal layer was added into a mixture of 30 μL luminol (Bio-Rad, Hercules, CA), 1 μL horseradish peroxidase (Jackson Immunoresearch, West Grove, PA), and 69 μL Milli-Q water in each well of a 96-well plate. Chemiluminescence (RLU, relative luminescent units) was detected from the ROS signals after 5 min incubation using a GloMax^®^ 96 Microplate Luminometer (Promega, Madison, WI).

### Malondialdehyde (MDA) Assay

The malondialdehyde (MDA) assay was performed to determine lipid peroxidation in rice leaf sheath tissues as described previously (Zhang et al., 2009; Dangol et al., 2019). Lipid peroxidation is the degradation of lipids due to oxidative damage in plant cells. Briefly, the ground tissue powder of rice leaf sheath was mixed with the reaction solution [0.5% (w/v) thiobarbituric acid, 20% (v/v) trichloroacetic acid (TCA), and 0.25 mL 175 mM NaCl in 2 mL of 50 mM Tris-Cl, pH 8.0]. The mixed reaction was then incubated in boiling water for 5 min, cooled on ice for 5 min, and centrifuged at 14,000× *g* for 5 min. The MDA concentration (C) in the resulting supernatant was determined by measuring supernatant absorbances (OD, optical density) at 450, 532, and 600 nm, and then calculating MDA concentration according to the following equation: C = 6.45 × (OD_532_–OD_600_) – (0.56 × OD_450_).

### Ferric Ion (Fe^3+^) Detection by Prussian Blue Staining

Prussian blue staining was performed to detect ferric ion (Fe^3+^) in rice leaf sheaths as described previously (Liu et al., 2007; Dangol et al., 2019). Briefly, epidermal layers of rice leaf sheaths were excised and incubated in equal volumes (1:1, v/v) of 7% potassium ferrocyanide and 2% hydrochloric acid (HCl) for 15 h at room temperature. Prussian blue (ferric ferrocyanides, which combine with Fe^3+^ inside leaf sheath epidermal cells) was observed as a bright blue color under a fluorescence microscope. Prussian blue-stained cells were categorized into two phenotypes: Type I, cells that contain IH but are weakly or not Prussian blue-stained; and Type II, strongly Prussian blue-stained cells with only a few poor hyphae.

### Subcellular Localization of OsMEK2, OsMPK1, and OsWRKY90 in *N. benthamiana* Leaves

The full length cDNA of *OsMEK2*, *OsMPK1,* and *OsWRKY90* were amplified from the rice cDNA library with the gene-specific primers containing attB1 and attB2 sites, as described in Supplementary Table S1. The amplified PCR products were used as a template for the second PCR using attB1 and attB2 primers. The second PCR products were sub-cloned into the pDONR™201 entry vector using BP clonase (Invitrogen) to create entry clones. The entry clones were recombined into the Gateway binary vector pGWB552 tagged with G3 green fluorescent protein (G3GFP) using LR clonase (Invitrogen) (Nakagawa et al., 2007; Dangol et al., 2017).

The binary plasmids containing *OsMEK2*, *OsMPK1,* and *OsWRKY90* were transformed into *A. tumefaciens* GV3101. Recombinant agrobacteria were prepared for infiltration using a protocol as described previously with slight modification (Sainsbury and Lomonossoff, 2008). Briefly, single colonies of recombinant agrobacteria were incubated into the liquid LB media (10 g/L tryptone, 5 g/L yeast extract; 10 g/L NaCl, pH 7) containing spectinomycin (100 μg/L) for overnight at 28 °C with continuous shaking. Harvested recombinant agrobacteria were resuspended to an OD_600_ = 0.2 in MMA (10 mM MES pH 5.6, 10 mM MgCl_2_, 150 μM acetosyringone). The agrobacterial suspension was incubated for 2 h at room temperature, and infiltrated into the abaxial leaves of 6-week-old *Nicotiana benthamiana* plants using a blunt tipped plastic syringe. Subcellular localization of 00:GFP, OsMEK2:GFP, OsMPK1:GFP, and OsWRKY90:GFP in *N. benthamiana* epidermal cells 36 h after agroinfiltration were microscopically observed following 4’,6-diamidino-2-phenylindole (DAPI, 5μg/ml) staining for 10 min. Nuclear localization of the proteins was visualized by counterstaining the nuclei of the cells with DAPI.

### Microscopy

Images were captured using a fluorescence microscope (Zeiss equipped with Axioplan 2; Campbell, CA) with 40× oil-immersion objective lens. CM-H_2_DCFDA-specific fluorescence was visualized under the fluorescence microscope using a combination of excitation (450‒490 nm) and emission (515‒565 nm) green fluorescence (GF) filters. Subcellular images were also taken using a fluorescence microscope (Olympus, Japan) using bright field, GF (Ex/Em: 488/498–548 nm), and DAPI (Ex/Em: 405/421-523 nm) filters.

### Accession Numbers

Sequence data from this article can be found in the GenBank/EMBL data libraries under the following accession numbers: *OsMEK1* (Os01g32660), *OsMEK2* (Os06g05520), *OsMEK3* (Os03g12390), *OsMEK4* (Os02g46760), *OsMEK5* (Os06g09190), *OsMEK6* (Os02g54600), *OsMEK7* (Os06g09180), *OsMEK8* (Os06g27890), *OsMPK1* (Os06g06090), *OsMPK6* (Os10g38950), *OsWNK1* (Os07g38530), *OsWRKY90* (Os09g30400), *OsNADP-ME* (Os01g52500), *OsRbohB* (Os01g25820), *OsPR-1b* (Os01g28450), *OsPAL1* (Os04g43760), *OsAPX1* (*Os0.g17690*), *OsAPX2* (*Os07g49400*), *OsUbiqutin* (Os06g46770), *18S rRNA* (XR_003238819.1), *AtMKK1* (At4g26070), *AtMKK2* (At4g29810), *AtMEK3* (NP_198860), *AtMEK4* (At1g51660), *AtMEK5* (At3g21220), *AtMKK6* (At5g56580), *AtMKK9* (At1g73500), *AtWNK9* (At3g04910), *NtNPK2* (BAA06731), *SlMKK1* (NP_001234744), *SlMKK2* (NP_001234588), *NbMEK2* (LOC107818847), and *NtMEK2* (AF325168).

## Supplemental Data

The following materials are available in the online version of this article.

**Supplemental Figure S1.** Amino acid sequence alignment of rice MAPKKs (OsMEKs) with Arabidopsis, tomato, and tobacco MEKs.

**Supplemental Figure S2.** Circular phylogenetic tree of rice MAPKKs (OsMEKs) with Arabidopsis, tomato, and tobacco MAPKs.

**Supplemental Figure S3.** Nucleotide sequences and deduced amino acid sequences of rice MAP kinase kinase 2 (OsMEK2) genomic DNA.

**Supplemental Figure S4.** Quantitative real-time RT-PCR analysis of time-course expression of defense-response genes *OsPR-1b*, *OsPAL1*, *OsAPX1*, and *OsAPX2* in wild-type (WT) rice (cultivar DJ) and *ΔOsmek2* mutant during *Magnaporthe oryzae* 007 infection.

**Supplemental Figure S5.** Images of Prussian blue-stained and DAB-stained leaf sheath cells of wild-type (WT) rice (cultivar DJ) at different times after erastin treatment.

**Supplemental Figure S6.** ROS accumulation and lipid Peroxidation in leaf sheaths of wild-type (WT) rice (cultivar DJ) and *ΔOsmek2* mutant at 72 and 96 h after inoculation with avirulent *Magnaporthe oryzae* 007 in 10 μM erastin.

**Supplemental Figure S7.** Transcriptional analysis of *OsMPK1* expression in leaf sheath epidermal cells of wild-type (WT) rice (cultivar NB) and 35S:*OsMPK1* mutants using qRT-PCR.

**Supplemental Table S1.** Primers used in this study.

## SUPPLEMENTAL DATA

**Supplemental Figure S1. Amino acid sequence alignment of rice MAPKKs (OsMEKs) with Arabidopsis, tomato, and tobacco MEKs.**

Amino acid sequence of OsMEK2 was aligned with MAP kinase kinases of Arabidopsis, tomato, and tobacco using CLUSTAL OMEGA (EMBL-EBI). Accession numbers of the aligned plant MAPKKs are OsMEK1 (Os01g32660), OsMEK2 (Os06g05520), OsMEK3 (Os03g12390), OsMEK6 (Os02g54600), OsMEK8 (Os06g27890), AtMKK1 (At4g26070), AtMKK2 (At4g29810), AtMEK3 (NP_198860), AtMEK4 (At1g51660), AtMEK5 (At3g21220), SlMKK2 (NP_001234588), NtMEK2 (AF325168), and NbMEK2 (LOC107818847). Asterisks and dots at the bottom of sequences indicate identical and similar amino acids, respectively. Domain numbers (I~XI) on the top of sequences indicate conserved subdomains. The conserved consensus motif (GXGXXG) in conserved subdomain I is boxed. The active motif [D(I/L/V)K] and the conserved consensus motif [S/TXXXXXS/T, serine(S)/threonine (T) residues] between conserved subdomains VII and VIII are shown in yellow.

**Supplemental Figure S2. Circular phylogenetic tree of rice MAPKKs (OsMEKs) with Arabidopsis, tomato, and tobacco MAPKs.**

Phylogenetic tree was constructed using the neighbor-joining method based on Molecular Evolutionary Genetics Analysis Version 7.0 (MEGA7) (Kumar et al., 2016). Accession numbers of the plant MAPKKs are OsMEK1 (Os01g32660), OsMEK2 (Os06g05520), OsMEK3 (Os03g12390), OsMPK8a (Os06g27890), AtMEK3 (NP_198860), AtMEK4 (At1g51660), AtMEK5 (At3g21220), AtMKK1 (At4g26070), AtMKK2 (At4g29810), SlMKK1 (NP_001234744), SlMKK2 (NP_001234588), NtMEK2 (AF325168), NtNPK2 (BAA06731), and NbMEK2 (LOC107818847).

**Supplemental Figure S3. Nucleotide sequences and deduced amino acid sequences of rice MAP kinase kinase 2 (OsMEK2) genomic DNA.**

Small letters represent nucleotide sequences of exons and introns. Capital letters represent deduced amino acid sequences of exons. Initiation and termination codons of the *OsMEK2* coding region are represented by asterisks. Exon-intron splice junctions (gt/ag) are represented by bold letters. Numbers at the right refer to nucleotide and amino acid (in parentheses) residue positions in the respective sequence.

**Supplemental Figure S4. Quantitative real-time RT-PCR analysis of time-course expression of defense-response genes***OsPR-1b*, *OsPAL1*, *OsAPX1***, and***OsAPX2* **in wild-type (WT) rice (cultivar DJ),***ΔOsmek2,* **and 35S:***OsMEK2* **mutant plants during***Magnaporthe oryzae* **007 infection.**

Leaf sheaths of wild-type (cultivar DJ), *ΔOsmek2,* and 35S:*OsMEK2* mutant rice plants were sampled at different time points after inoculation, followed by total RNA extraction. Relative gene expression levels of defense-responsive genes (*OsPR-1b*, *OsPAL1*, *OsAPX1*, and *OsAPX2*) at each time point were obtained by normalizing the gene expression with respect to expression of the internal control 18S rRNA gene. Data represent the means ± SD from three independent experiments. Different letters above the bars indicate significantly different means (*P*<0.05) as analyzed by Fisher’s protected LSD test.

**Supplemental Figure S5. Images of Prussian blue-stained and DAB-stained leaf sheath cells of wild-type (WT) rice (cultivar DJ) at different time points after erastin treatment.**

Rice DJ leaf sheaths were treated with 10 μM erastin and water (Mock). Images were taken using a fluorescence microscope (Zeiss equipped with Axioplan 2). Prussian blue and DAB staining in erastin-treated rice leaf sheaths did not detect ferric ion and ROS accumulation, respectively, at different time points after treatment. Experiments were repeated three times with similar results. hpt, hours post-treatment. Scale bars=20 μm.

**Supplemental Figure S6. ROS accumulation and lipid peroxidation in leaf sheaths of wild-type (WT) rice (cultivar DJ) and***ΔOsmek2* **mutant plants at 72 and 96 h after inoculation with avirulent***Magnaporthe oryzae* **007 with 10 μM erastin.**

Rice *ΔOsmek2* mutant leaf sheaths were inoculated with conidial suspensions (4×10^5^ conidia mL^−1^) of avirulent *M. oryzae* 007 with and without 10 μM erastin. Avirulent *M. oryzae* 007 infection induced ROS production and lipid peroxidation in leaf sheaths of the susceptible *ΔOsmek2* mutant at 96 hpi, which was similar to those in erastin-treated leaf sheaths of the susceptible *ΔOsmek2* mutant. A, DAB-stained cell phenotypes at 72 and 96 h after inoculation. DAB-stained cells were categorized into two phenotypes: Type I, cells that contain invasive hyphae (IH) but are weakly or not DAB-stained; and Type II, strongly DAB-stained cells with only a few poor hyphae. Scale bars=20 μm. B, Quantification of DAB-stained cells at 72 and 96 h after inoculation. DAB-stained cells were categorized into two phenotypes: Type I, infected cells that display no or weak DAB staining; Type II, infected cells that display strong DAB staining. C, Determination of lipid (MDA) peroxidation in rice leaf sheaths at 72 and 96 h after inoculation. Results are presented as mean values ± SD; *n*=4 leaf sheaths from different plants. Images were captured using a fluorescence microscope (Zeiss equipped with Axioplan 2). Results are presented as mean values ± SD; *n*=4 leaf sheaths from different plants. Different letters above the bars indicate significantly different means (*P*<0.05) as analyzed by Fisher’s protected LSD test. Experiments were repeated three times with similar results. hpi, hours post-inoculation.

**Supplemental Figure S7. Transcriptional analysis of***OsMPK1* **expression in leaf sheath epidermal cells of wild-type (WT) rice (cultivar NB) and 35S:***OsMPK1* **mutants using qRT-PCR.**

Relative gene expression of *OsMPK1* was obtained by normalizing with respect to the expression of the internal control *OsUbiquitin* gene. Data represent the means ± SD from three independent experiments. Asterisks indicate statistically significant differences (Student’s *t* test, *P<* 0.01).

